# Microtubule Nucleation Promoters Mto1 and Mto2 Regulate Cytokinesis in Fission Yeast

**DOI:** 10.1101/843110

**Authors:** Samantha E. R. Dundon, Thomas D. Pollard

**Author notes:** Available as a preprint on BioRχiv: https://www.biorxiv.org/content/10.1101/843110v1 doi: https://doi.org/10.1101/843110.

## Abstract

Microtubules of the mitotic spindle direct cytokinesis in metazoans but this has not been documented in fungi. We report evidence that astral microtubules help coordinate cytokinetic furrow formation in fission yeast. The temperature-sensitive *cps1-191* strain (Liu et al., 1999) with a D277N substitution in β-glucan synthase 1 (Cps1/Bgs1) was reported to arrest with an unconstricted contractile ring. We discovered that contractile rings in *cps1-191* cells do constrict slowly and that an S338N mutation in the *mto2* gene is required with the *bgs1*_*D277N*_ mutation to reproduce the *cps1-191* phenotype. Complexes of Mto2 and Mto1 with γ-tubulin regulate microtubule assembly. Deletion of Mto1 along with the *bgs1*_*D277N*_ mutation also gives the *cps1-191* phenotype, which is not observed in *mto2*_*S338N*_ or *mto1Δ* cells expressing *bgs1+*. Both *mto2*_*S338N*_ and *mto1Δ* cells nucleate fewer astral microtubules than normal and have higher levels of Rho1-GTP at the division site than wild-type cells. We report multiple conditions that sensitize *mto1Δ* and *mto2*_*S338N*_ cells to furrow ingression phenotypes.

**Summary:** Dundon and Pollard show that compromising the Mto1 or Mto2 regulators of the fission yeast γ-tubulin complex reduces or eliminates astral microtubules, exaggerates the effects of a D277N substitution in β-glucan synthase 1 (Cps1/Bgs1) on the rate of cytokinetic furrow formation, and increases Rho1-GTP at the cleavage site.

## Introduction

Animals, fungi, and amoebas use an actomyosin contractile ring to divide. This ring is constructed at the end of the cell cycle in uninucleate cells, and cellular division must only proceed after successful DNA segregation to generate viable daughters. However, the mechanisms communicating DNA status to the contractile ring are still incompletely understood. Failure to correctly orchestrate this process can cause chromosome segregation errors and aneuploidy, which can promote chromosomal instability.

The fission yeast *Schizosaccharomyces pombe* is an excellent model for understanding mitosis and cytokinesis, including the most comprehensive cytokinetic parts list. The timing of mitotic and cytokinetic milestones is highly reproducible and well characterized; the stereotypical size and shape of *S. pombe* cells facilitates rigorous quantitative microscopy; and numerous temperature-sensitive alleles and facile genetic methods enable dissection of molecular processes.

*S. pombe* cells assemble a contractile ring from precursors called nodes over 10-15 min following separation of the spindle pole bodies (SPBs) at the onset of anaphase at 23°C (Vavylonis et al., 2008). During anaphase B, the ring matures by adding proteins such as the myosin-II isoform, Myp2 (Wu et al., 2003), the major motor for constriction (Laplante et al., 2015). The maturation period ends with onset of constriction 35 min after SPB separation, even if mutations slow the assembly of the ring (Tebbs and Pollard, 2013; Coffman et al., 2009; Roberts-Galbraith et al., 2010; Wang et al., 2014; Laplante et al., 2015). If a ring takes longer than 35 min to assemble, it constricts immediately (Chen and Pollard, 2011; Tebbs and Pollard, 2013; Li et al., 2016).

Three conditions appear to be required to begin furrow ingression: an intact contractile ring; a signal originating from the cell cycle clock; and initiation of septum synthesis. The enzyme β-glucan synthase 1 (Bgs1) is recruited with its regulator Rho1 to the equator where it synthesizes the primary septum (Arellano et al., 1996) and helps anchor the ring to generate inwardly directed force (Arasada and Pollard, 2014; Davidson et al., 2015). Furrow ingression initially depends on both tension produced by the contractile ring and septum deposition by Bgs1 (Zhou et al., 2015; Thiyagarajan et al., 2015), but after ~50% ingression, septum synthesis alone suffices to complete division (Proctor et al., 2012). Although the septum can be completed in the absence of the actomyosin ring, tension in the ring promotes septum deposition and maintains the circularity of the pore during septum deposition (Zhou et al., 2015; Thiyagarajan et al., 2015). Mutations compromising the contractility or tension in the ring decrease the constriction rate (Pelham and Chang, 2002; Proctor et al., 2012; Laplante et al., 2015; Tebbs and Pollard, 2013; Li et al., 2016). Ring constriction is faster if the internal pressure against which the furrow ingresses is reduced through mutation or increasing the osmolarity of the growth medium (Morris et al., 2019; Proctor et al., 2012). Bgs1, activated by Rho1, synthesizes the primary septum (Cortés et al., 2007). This is followed by synthesis of the secondary septum by Bgs4 (Rho1) and Ags1 (Rho2) (Cortés et al., 2012; 2005; Arellano et al., 1996; Calonge et al., 2000). While Rho1 activity is required for cytokinesis, it must decrease after septation to allow cell separation (Nakano et al., 1997).

The fascinating fission yeast strain *cps1-191* (Liu et al., 1999) attracted our attention, because it seemed to lack some aspect of the signal for initiating ring constriction. Liu et al. created this strain by chemical mutagenesis and reported that the cells arrested at 36°C with two nuclei and an unconstricted cytokinetic ring. They concluded that colonies did not grow at 36°C due to failed cytokinesis. They identified a candidate gene by genetic mapping, complemented temperature sensitivity of the strain with the wild type *cps1+/bgs1+* gene, identified the D277N substitution in the gene and named the strain *cps1-191* (Liu et al., 1999). Time-lapse microscopy of the *cps1-191* strain confirmed that the nuclei separate normally but actomyosin rings remain intact without constricting for an hour after assembly at the restrictive temperature (Arasada and Pollard, 2014). The *cps1-191* strain has been used in many studies to generate cells with non-constricting actomyosin rings (Pardo and Nurse, 2003; Yamashita et al., 2005; Venkatram et al., 2005; Wachtler et al., 2006; Loo and Balasubramanian, 2008; Roberts-Galbraith et al., 2009) and others.

Based on these reports, we selected the *cps1-191* strain to investigate the mechanisms that initiate cytokinetic ring constriction. However, we determine that rings in *cps1-191* cells do not arrest *per se*, but constrict very slowly and that cells with the *bgs1*_*D227N*_ mutation die from lysis at 36°C rather than arresting the cell cycle as previously thought. Surprisingly, we found that the *cps1-191* constriction phenotype depends on a second point mutation in the gene for the γ-tubulin regulator Mto2, implicating microtubules in the process that drives furrow ingression.

*S. pombe* has multiple microtubule organizing centers (MTOCs) containing γ-tubulin in addition to the SPB (Sawin and Tran, 2006). During interphase, multiple MTOCs localize along microtubule bundles (Janson et al., 2005). During mitosis, interphase microtubules disassemble and the SPBs nucleate spindle and astral microtubules. Finally, during cytokinesis Myp2 recruits MTOCs to the equator (Samejima et al., 2010), where they nucleate the post-anaphase array (PAA) of microtubules. Mto1 and, to a lesser extent, Mto2 are critical proteins for the function and cell-cycle regulated activities of these MTOCs. In the absence of Mto1, no MTOCs form beyond the SPB, which only nucleates microtubules from its inner face (Zimmerman and Chang, 2005). In biochemical reconstitution experiments, the Mto1/Mto2 complex and MOZART1 homolog Mzt1 are required for γ-tubulin to promote microtubule nucleation (Leong et al., 2019). We find that Mto1/Mto2 complex mutants nucleate fewer than normal numbers of astral microtubules, and report one consequence, higher than normal Rho1-GTP activity at the cleavage site. Our findings reveal a new link between microtubules and furrow ingression in fission yeast.

## Results

### Cytokinetic furrows ingress slower in the *cps1-191* strain than in cells with the *bgs1_D277N_* mutation

Long time-lapse movies of the *cps1-191* strain with genome-encoded Rlc1-tdTomato (regulatory light chain for both isoforms of myosin-II, Myo2 and Myp2) revealed that the actomyosin ring constricted ~30-fold slower (median 0.02 μm/min) at the restrictive temperature of 36°C than in wild-type cells (median 0.62 μm/min) (Fig. 1A, B). We never observed abnormal ring structures such as detachment from the plasma membrane (Arasada and Pollard, 2014; Laplante et al., 2015; Cheffings et al., 2019), therefore we consider Rlc1-tdTomato constriction to reflect furrow ingression. Thirty percent of these fully-formed contractile rings slid along the membrane of *cps1-191* cells at 36°C, as reported (Arasada and Pollard, 2014; Cortés et al., 2015). We shifted *cps1-191* cells from the permissive (25°C) to restrictive (36°C) temperature on the microscope to determine the time required for the phenotype to manifest. When the SPBs separated within 30 min of the temperature shift, furrow ingression was normal. If SPB separation occurred >30 min after the shift, furrows did not form normally (Fig S1A).

**Figure 1.**
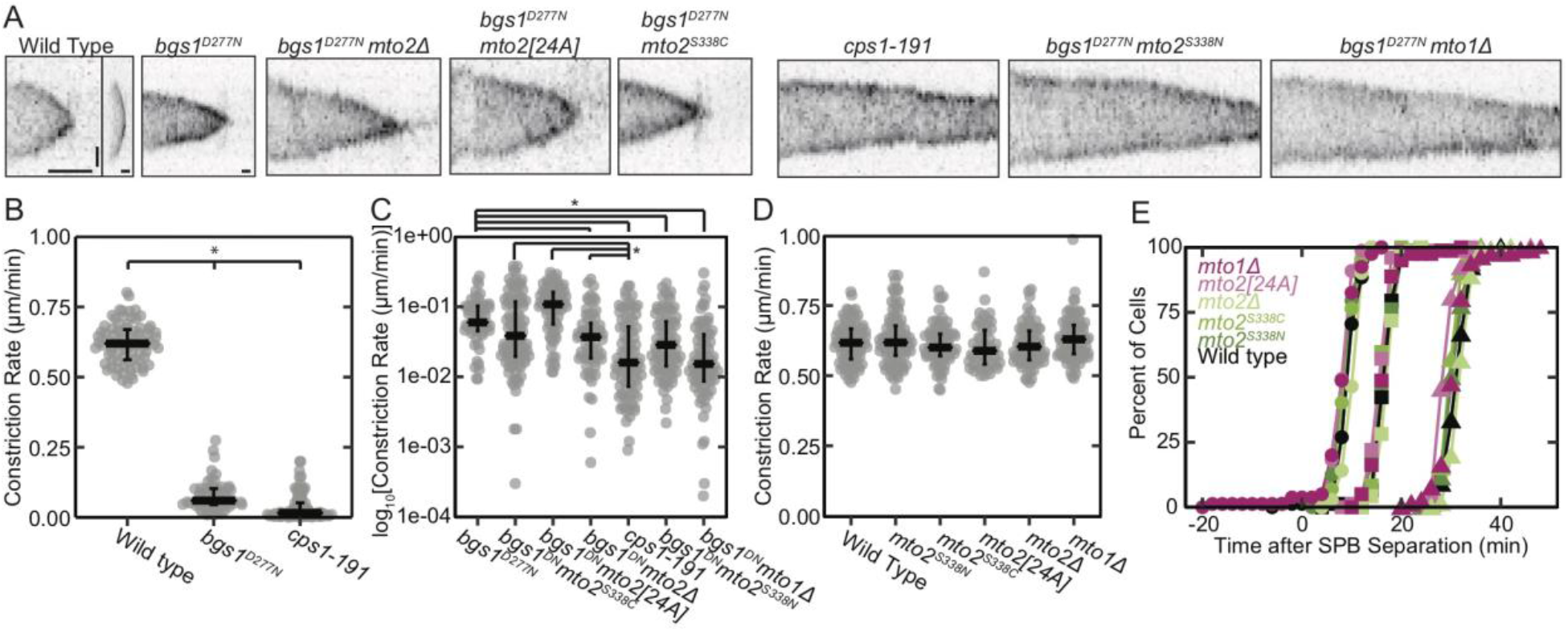
Both the *bgs1*_*D277N*_ and the *mto2*_*S338N*_ mutations are required to cause the *cps1-191* constriction phenotype in a wild-type background. A) Kymographs of inverted contrast, maximum-intensity projected images of contractile rings in strains with Rlc1-tdTomato at 36°C. Wild-type cells were imaged at 1-min intervals, and *bgs1*_*D277N*_ and *cps1-191* cells were imaged with 5-min intervals. The kymograph of the wild-type cell is displayed (left sub-panel) as acquired and (right sub-panel) rescaled to match the timescale of the kymographs (other panels) of the *cps1-191* and six different *bgs1*_*D277N*_ strains. Horizontal scale bars = 15 min, vertical scale bar = 1 μm. B) Rates of cytokinetic ring constriction measured from a subset of kymographs in A. The data are not normally distributed, so the median and 1_st_ and 3_rd_ quartiles are indicated by black bars, n ≥ 55 cells. C) Log_10_-transformed cytokinetic ring constriction rates of cells carrying the *bgs1*_*D277N*_ mutation measured from kymographs as in A. The median and 1_st_ and 3_rd_ quartiles are indicated by black bars, n ≥ 57 cells. Significance was determined by Welch’s ANOVA followed by Tukey post-hoc test (p < 0.05). D) Cytokinetic ring constriction rates of cells carrying *bgs1+* measured from kymographs as in A. The median and 1_st_ and 3_rd_ quartiles are indicated by black bars. No significant differences were detected by Welch’s ANOVA. E) Cumulative distribution plots showing accumulation of cells with rings that have (•) assembled, (■) initiated constriction, and (▲) completed constriction in wild-type and *bgs1+* cells with various additional mutations at 36°C. n ≥ 71 cells for C and D.

We made a strain with just the *bgs1*_*D277N*_ mutation in a wild-type background and found that furrowing progressed 3-fold faster (median 0.06 μm/min) than in *cps1-191* cells. Nevertheless, these *bgs1*_*D277N*_ cells had a growth defect at 36°C similar to the *cps1-191* strain in serial dilution assays, consistent with the 2:2 segregation previously observed for this phenotype (Fig. S1B) (Liu, et al., 1999). Both *bgs1*_*D277N*_ and *cps1-191* cells lysed frequently during imaging (Fig. S1D), which caused the growth defect at 36°C on solid medium. None of the *bgs1+* strains we examined lysed during observations. The lysis frequency varied substantially between replicate samples, suggesting that this phenotype is extremely sensitive to minute environmental differences. Consistent with this hypothesis, the osmotic stabilizer sorbitol partially rescued the growth of *cps1-191* and *bgs1*_*D277N*_ cells at 36°C (Fig. S1C).

### The *cps1-191* strain carries a large number of mutations

We sequenced the complete genome of the *cps1-191* strain. Comparison to the *S. pombe* reference genome obtained from PomBase (Wood et al., 2002; Lock et al., 2019) revealed 384 unique mutations, 271 of which were transition mutations consistent with the nitrosoguanidine mutagenesis used to generate the strain (Balasubramanian et al., 1998) (Fig. S1E). Ten of these transition mutations substituted amino acids in eight genes located across all three chromosomes, including the reported *bgs1*_*D277N*_ mutation (Fig. S1F, Table 1).

**Table 1.**
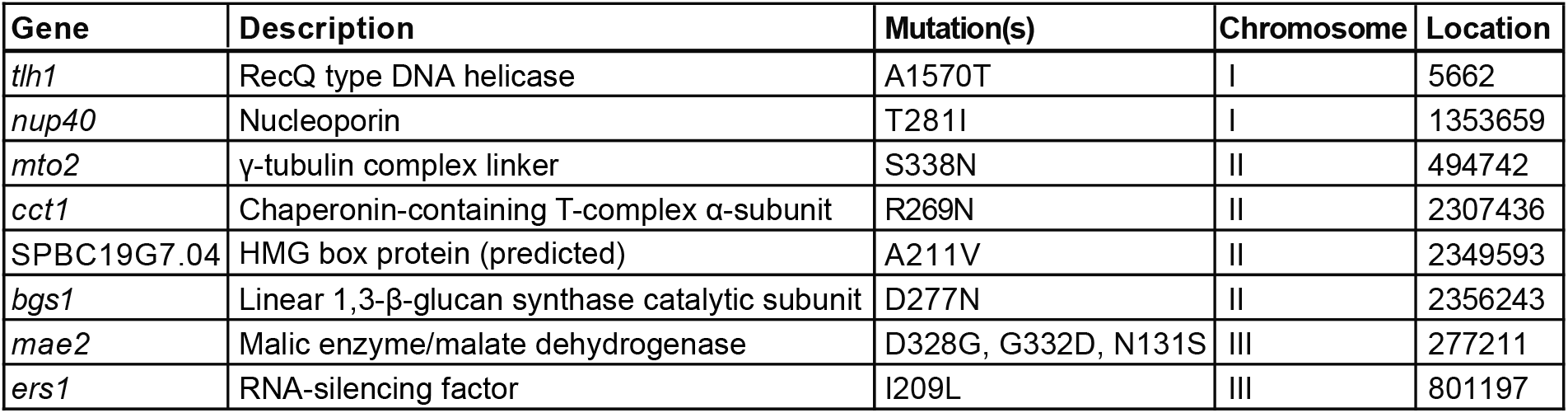
Mutations in ORFs in the *cps1-191* strain.

| Gene | Description | Mutation(s) | Chromosome | Location |
| --- | --- | --- | --- | --- |
| *tlh1* | RecQ type DNA helicase | A1570T | I | 5662 |
| *nup40* | Nucleoporin | T281I | I | 1353659 |
| *mto2* | γ-tubulin complex linker | S338N | II | 494742 |
| *cct1* | Chaperonin-containing T-complex α-subunit | R269N | II | 2307436 |
| SPBC19G7.04 | HMG box protein (predicted) | A211V | II | 2349593 |
| *bgs1* | Linear 1,3-β-glucan synthase catalytic subunit | D277N | II | 2356243 |
| *mae2* | Malic enzyme/malate dehydrogenase | D328G, G332D, N131S | III | 277211 |
| *ers1* | RNA-silencing factor | I209L | III | 801197 |

### Mutations in the Mto1/Mto2 complex synthetically interact with the *bgs1_D277N_* mutation

To determine which mutations contribute to the *cps1-191* phenotype, we examined the PomBase annotations for each protein-coding gene with a substitution mutation (Table 1), looking for roles in cell cycle regulation or cytokinesis (Lock et al., 2019). Two of the eight affected genes, Mto2 and Nup40, met this criterion. To determine if either or both of these mutations affected the rate of furrow ingression in combination with *bgs1*_*D277N*_, we took advantage of the fact that we saved multiple progeny from crosses generating *cps1-191* strains with labeled rings and SPBs. These strains were already confirmed to contain the *bgs1*_*D277N*_ mutation by sequence analysis. We measured the furrowing rates at 36°C and determined the *mto2* and *nup40* mutation status for each of these strains. Only presence of the *mto2*_*S338N*_ mutation correlated with the slowest furrowing rate observed for *cps1-191* (Table S1). Double mutants bearing *bgs1*_*D277N*_ *nup40*_*T281I*_ exhibited ingression rates no different from *bgs1*_*D277N*_ alone. The *mto2*_*S338N*_ mutation was also particularly interesting due to previous work linking Mto2 to the cytokinetic ring (Samejima et al., 2010).

Combining the *bgs1*_*D277N*_ and *mto2*_*S338N*_ mutations in a wild-type background reproduced the slow constriction rate of *cps1-191* (Fig. 1A, C). The Mto2 C terminus is highly phosphorylated (Borek et al., 2015), so we tested if the absence of phosphorylation at S338 caused the synthetic interaction with *bgs1*_*D277N*_. Neither S338C, which precludes phosphorylation of S338, nor the [24A] strain with 24 phosphorylated serines mutated to alanines (Borek et al., 2015), constricted as slowly as the *bgs1*_*D277N*_ *mto2*_*S338N*_ or *cps1-191* mutants, but instead resembled the *bgs1*_*D277N*_ strain with a single mutation (Fig. 1A, C).

Mto2 contributes to the formation and regulation of MTOCs other than SPBs in *S. pombe* through interactions with its binding partner Mto1 (Sawin and Tran, 2006). Neither *mto1+* nor *mto2+* are essential genes, but mutations of *mto1+* are more severe than mutations of *mto2+* (Sawin et al., 2004; Venkatram et al., 2005). In a *bgs1+* background with Rlc1-tdTomato to label contractile rings and Pcp1-mEGFP to label SPBs, all mutant strains of *mto1* and *mto2* ingressed furrows at the wild-type rate with cytokinesis timelines similar to wild-type cells (Fig. 1D, E). In a *bgs1*_*D277N*_ strain deletion of Mto1 slowed furrowing to the rate observed in *cps1-191* cells, while *bgs1*_*D277N*_ *mto2Δ* cells exhibited an intermediate phenotype (Fig. 1C). Therefore, the *bgs1*_*D277N*_ mutation sensitizes cells to a compromised Mto1/Mto2 complex.

We hypothesized that recruitment of Bgs1 to the division site may be defective in *mto1Δ* and *mto2*_*S338N*_ cells, increasing their sensitivity to loss of Bgs1 function. However, *bgs1+mto1Δ* and *bgs1+mto2*_*S338N*_ cells recruited the same number of the number of Bgs1 molecules recruited to the equator during cytokinesis as wild-type cells (Fig. S1G).

### Mitotic microtubules are disrupted in *mto1Δ* and *mto2_S338N_* cells

We investigated the microtubule cytoskeleton in *mto1Δ* and *mto2*_*S338N*_ cells to determine common phenotypes that may affect furrow ingression. Over the course of spindle elongation in wild-type cells each SPB nucleated multiple, short-lived astral microtubules with a median duration of 20 s (n = 175 microtubules) (Fig. 2A, S2A). The *mto2*_*S338N*_ strain produced fewer astral microtubules than normal, and most SPBs in *mto1Δ* cells nucleated no astral microtubules, as reported (Sawin et al., 2004)(Fig. 2B, C). We measured the numbers of microtubules nucleated by SPBs only during anaphase B, to avoid intra-nuclear “astral” microtubules nucleated prior to this stage, which may not be affected by *mto1* loss (Zimmerman et al., 2004).

**Figure 2.**
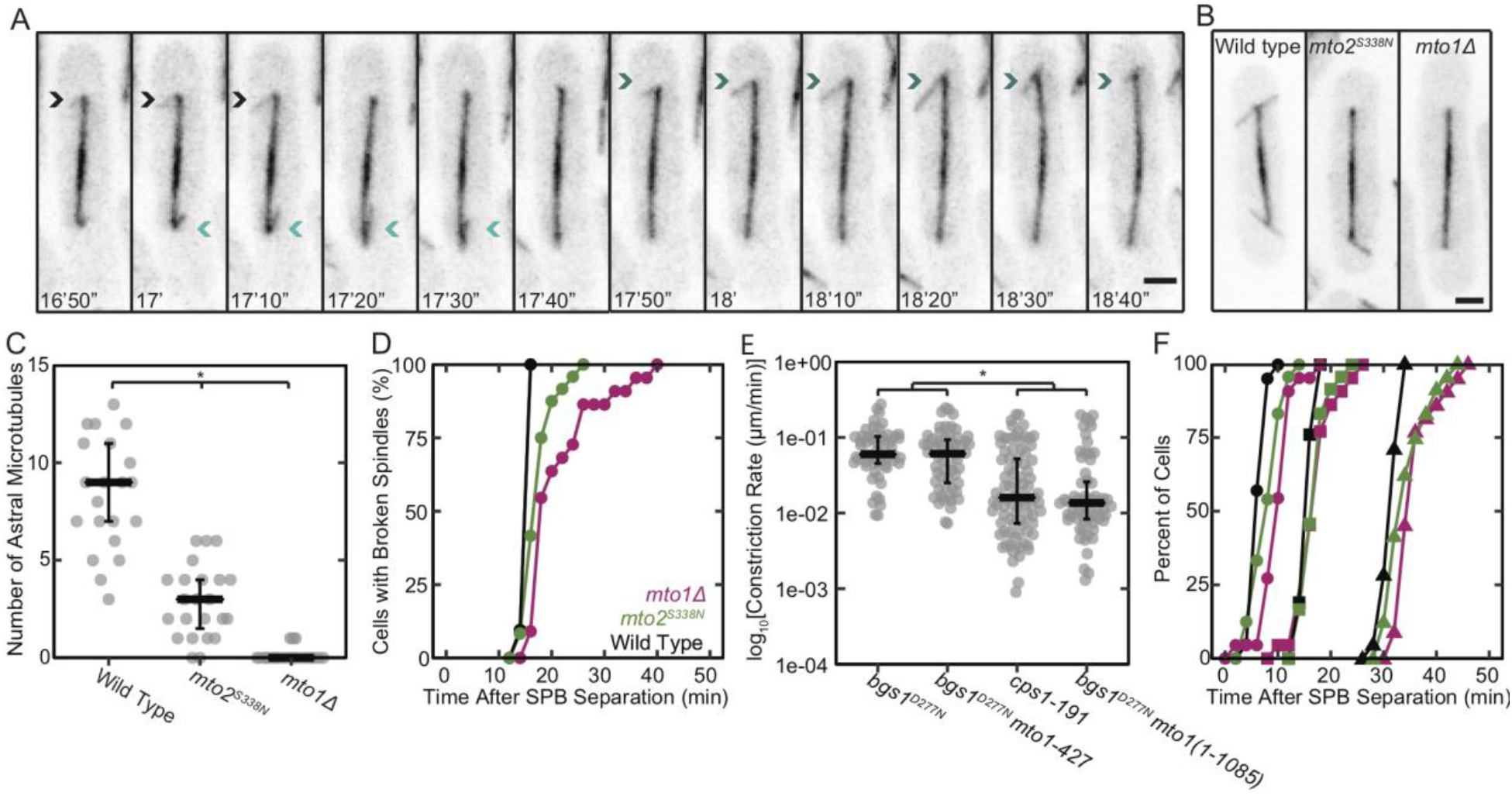
The *mto2*_*S338N*_ and *mto1Δ* mutations perturb astral microtubules and the time of spindle breakage. A) Time series of inverted contrast maximum-intensity projected fluorescence micrographs of wild type cells at 36°C with astral microtubules labeled with GFP-Atb2 (α tubulin). Arrows of the same color mark the same astral microtubule in consecutive frames. Scale bar 2 μm. B) Representative fluorescence micrographs of three strains (wild type, *mto2*_*S338N*_, and *mto1Δ*) at 36°C with spindle microtubules labeled with GFP-Atb2. Scale bar 2 μm. C) The number of astral microtubules nucleated from a single SPB over the course of mitosis at 36°C in wild-type, *mto2*_*S338N*_, and *mto1Δ* cells, n ≥ 21 spindle poles. Significance was determined using pairwise K-S tests with Bonferroni correction. D & F) Cumulative distribution plots comparing mitotic and cytokinetic outcomes in wild type, *mto2*_*S338N*_, and *mto1Δ* cells with GFP-Atb2 to mark microtubules, Rlc1-tdTomato to mark the cytokinetic ring, and Sfi1-mCherry to mark SPBs at 36°C, n ≥ 21 cells. D) Time course after time zero of the accumulation of cells with broken mitotic spindles. E) Log_10_-transformed cytokinetic ring constriction rates in strains with Rlc1-tdTomato at 36°C lacking either the PAA microtubules (*mto1-427*) or astral microtubules (*mto1(1-1085)*) in combination with *bgs1*_*D277N*_. The median and 1_st_ and 3_rd_ quartiles are indicated by black bars, n ≥ 57 cells. Significance was determined by Welch’s ANOVA followed by Tukey post-hoc test (p < 0.05). F) Time course after time zero of the accumulation of cells with rings that have (•) assembled, (■) initiated constriction, and (▲) completed constriction.

Spindle breakage, the time at which the spindle began to disassemble after reaching its maximum length, occurred later than normal in both the *mto2*_*S338N*_ and *mto1Δ* strains (Fig. 2D, p < 0.05 by pairwise K-S tests with Bonferroni correction). Both spindle breakage and formation of the PAA occur during the ring maturation stage in wild-type cells (Fig. 2D, S2C).

The PAA is nucleated from γ-tubulin complexes associated with the cytokinetic ring (Samejima et al., 2010). Therefore, we considered whether a disruption of the PAA microtubules nucleated by the Mto1/Mto2 complex causes the synthetic constriction phenotypes we observe. As reported (Sawin et al., 2004), we did not observe PAA formation in *mto1Δ* cells, but the PAA formed at the normal time in *mto2*_*S228N*_ cells (Fig. S2C). This suggested that the observed loss of astral microtubules and not loss of PAA microtubules causes a synthetic phenotype with *bgs1*_*D277N*_. To assess this hypothesis more definitively, we took advantage of the *mto1-427* mutant that lacks PAA microtubules, and the *mto1(1-1085)* mutant that lacks astral microtubules (Samejima et al., 2010). Only the *mto1(1-1085)* mutant exhibited the synthetic effect on constriction rate when combined with the bgs1D277N mutation (Fig. 2E). Therefore, loss of Mto1 from the outer face of the SPB and inability to nucleate astral microtubules synthetically interacts with the *bgs1*_*D277N*_ mutation. In contrast, selective loss of Mto1 from the division site and lack of PAA microtubules has no effect on the constriction rate in this background. It remains possible that a function of Mto1 and Mto2 independent of microtubule nucleation contributes to the synthetic interaction with *bgs1*_*D277N*_, but the cells repolymerize their microtubules within one hour of continuous exposure to even a high dose (250 μg/mL) of carbendazim, so insufficient time was available to measure furrow ingression in *bgs1*_*D277N*_ cells without microtubules.

Although the cytokinetic timelines of *mto2*_*S338N*_ and *mto1Δ* cells were indistinguishable from wild-type cells at 36°C (Fig. 1E), expression of GFP-tagged α-tubulin (Atb2) caused synthetic phenotypes with these mutations. Ring assembly was slightly delayed in *mto2*_*S338N*_ and *mto1Δ* cells carrying GFP-Atb2 compared with wild-type cells bearing GFP-Atb2 (Fig. 2F, p < 0.05 by pairwise K-S tests with Bonferroni correction, n ≥ 21 cells). Completion of ring constriction was also delayed in *mto1Δ* cells in this experiment. This substantiates that the *mto1Δ* and *mto2*_*S338N*_ mutations perturb cytokinesis when combined with additional stressors. We believe that the effects we observe on astral microtubule nucleation are not an artifact of GFP-Atb2 expression, as they are consistent with *mto1Δ* and *mto2Δ* microtubule-nucleation defects shown by immunofluorescence (Sawin et al., 2004; Janson et al., 2005; Samejima et al., 2005).

### Neither *mto1Δ* nor *mto2_S338N_* cells exhibit SIN phenotypes

We wondered whether the septation initiation network (SIN) signaling pathway operates normally in the *mto1Δ* and *mto2*_*S338N*_ mutants. The SIN (Hippo in humans and mitotic exit network (MEN) in budding yeast) promotes formation of the contractile ring and cytokinesis (Simanis, 2015). SIN components localize to the SPB (Simanis, 2015), and it is possible that the loss of astral microtubule nucleation capacity also affects SIN signaling. Hyperactivation of SIN promotes septum formation during interphase (Minet et al., 1979), while low SIN activity results in unstable contractile rings and unreliable cytokinesis, promoting formation of multinucleate cells (Mitchison and Nurse, 1985; Dey and Pollard, 2018). Cytokinetic rings in *cps1-191*, *bgs1*_*D277N*_, and *mto2*_*S338N*_ cells are so stable that we never observed one disassemble, suggesting that the SIN activity is normal.

We rarely observed the formation of binucleate interphase cells in the *mto1Δ* mutant cells (6 of 220 cells) and never in *mto2*_*S338N*_ cells (0 of 234 cells). The *mto1Δ* cells have defects in nuclear positioning (Zimmerman and Chang, 2005), so both nuclei occasionally migrate into one daughter compartment after the contractile ring forms, resulting in one binucleate and one anucleate daughter cell. Contractile rings in these cells formed at the correct time in mitosis and did not disassemble. Early in mitosis, the SIN kinase Cdc7 first localizes to both SPBs and then to only one SPB during anaphase B (Sohrmann et al., 1998). Cdc7 localizes to one SPB in *cps1-191* cells with an unconstricted ring, which suggests normal SIN activity (Liu et al., 1999). We measured the number of Cdc7-mEGFP molecules at each SPB in *mto2*_*S338N*_ and *mto1Δ* cells and did not detect a significant difference from the wild type in either the peak number of molecules or the timing of the peak after SPB separation (Fig S2D-G).

### *mto1Δ* modulates the effects of salt stress on cytokinesis

Acute salt stress produced by addition of 0.6 M KCl to rich medium (YE5S) causes a delay in ring constriction particularly in cells lacking *myp2*, which encodes one of two type-II myosins in fission yeast (Bezanilla et al., 1997; Okada et al., 2019). Myp2 recruits Mto1 to the ring and is required to generate the PAA (Samejima et al., 2010). We hypothesized that if loss of Mto1 at the equator causes sensitization to salt stress in *myp2Δ* cells, we would observe the same effect of salt stress in *mto1Δ* and *mto2*_*S338N*_ cells. We found that 0.6 M KCl also decreases the rate of ring constriction in minimal medium (EMM5S) in wild-type, *mto2*_*S338N*_, and *mto1Δ* cells (Fig. 3A). Surprisingly, rings in *mto1Δ* cells constricted faster than in wild-type cells in the presence of this stress, though still more slowly than in the absence of KCl, so rings completed constriction earlier than in the other two strains in the presence of KCl (Fig. 3C). However, the *mto1Δ* and *mto2*_*S338N*_ phenotypes are distinct from the *myp2Δ* phenotype under salt stress (Okada et al., 2019), so the loss-of-function phenotypes differ for the Mto1/Mto2 complex and Myp2.

**Figure 3.**
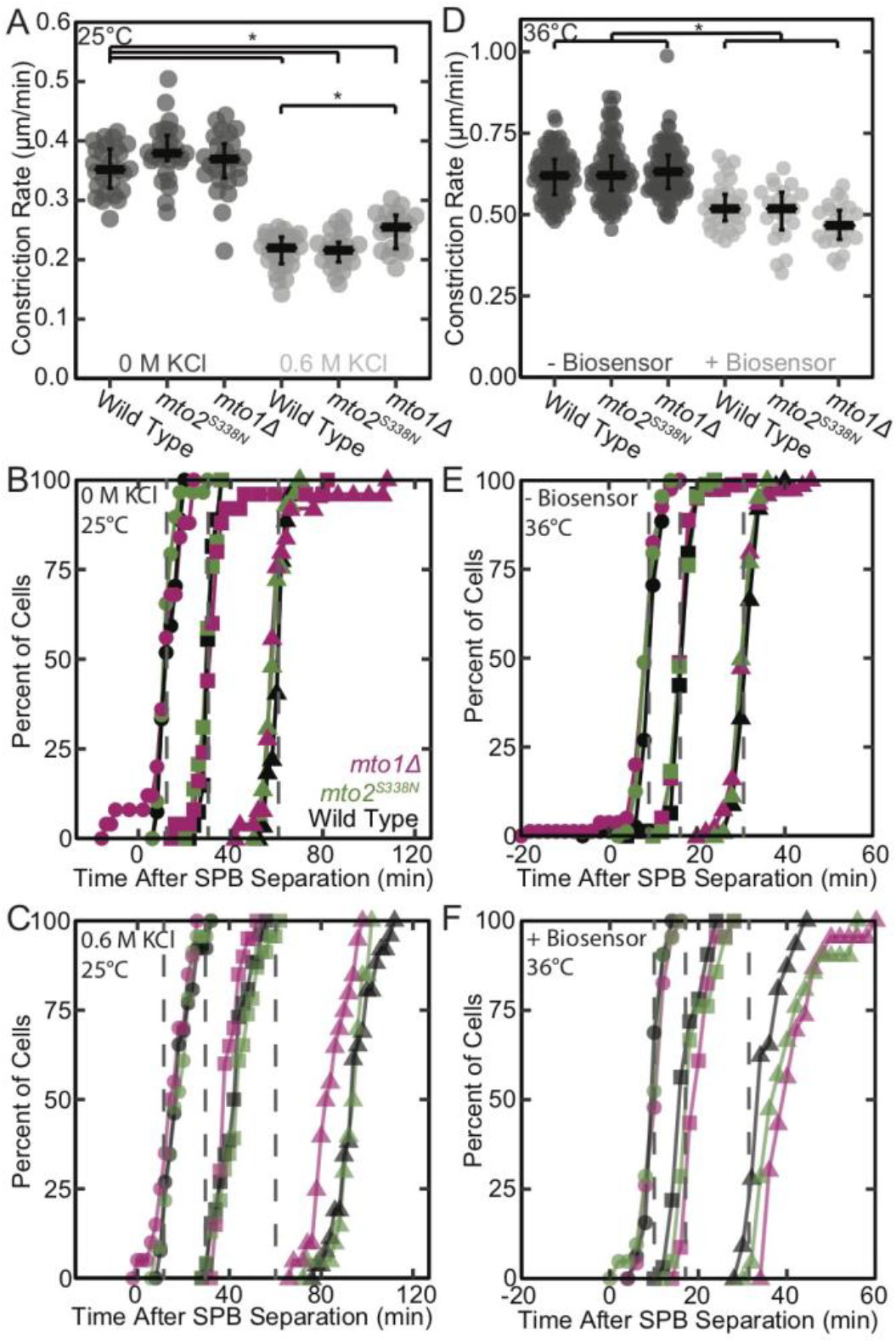
The *mto2*_*S338N*_ and *mto1Δ* mutations sensitize cells to stresses. A-C) Effects of 0.6 M KCl on cytokinesis in wild type, *mto2*_*S338N*_, and *mto1Δ* cells at 25°C. A) Rates of cytokinetic ring constriction. The median and 1_st_ and 3_rd_ quartiles are indicated by black bars and significance (p < 0.05) was determined using Welch’s ANOVA followed by Tukey post-hoc test. B & C) Time courses of cytokinetic events in cells expressing Rlc1-tdTomato to mark the cytokinetic ring and Pcp1-mEGFP to mark SPBs. Cumulative distribution plots show the accumulation of cells with rings that have (•) assembled, (■) initiated constriction, and (▲) completed constriction in medium with (B) no added KCl or (C) 0.6 M KCl. Dashed gray lines in (B) and (C) indicate the time when 50% of cells in the (B) reach each milestone to facilitate comparison. n ≥ 20 cells for A-C. D-F) Effects of the Rho1-GTP biosensor Pkc1(HR1-C2) on cytokinesis at 36°C. D) Rates of cytokinetic ring constriction. The median and 1_st_ and 3^rd^ quartiles are indicated by black bars and significance (p < 0.05) was determined using Welch’s ANOVA followed by Tukey post-hoc test. E & F) Cumulative distribution plots comparing mitotic and cytokinetic outcomes in cells (E) without and (F) with the Pkc1(HR1-C2) Rho1 biosensor. Plots show the accumulation of cells with rings that have with (•) assembled, (◼) initiated constriction, and (▲) completed constriction. Dashed gray lines in (E) and (F) indicate the time when 50% of cells in the (F) reach each milestone to facilitate comparison. n ≥ 21 cells for D-F.

### The *mto1Δ* and *mto2_S338N_* cells have high Rho1-GTP at the cleavage site

To explore one of many possible pathways influenced by the defects in astral microtubules, we localized *S. pombe* Rho1-GTP using the Pkc1(HR1-C2)-mEGFP biosensor (Davidson et al., 2015). This probe, derived from *Saccharomyces cerevisiae* Pkc1, contains Rho1-GTP binding HR1 domains and a plasma-membrane targeting C2 domain fused to mEGFP (Kono et al., 2012). When in the active, GTP-bound state, RhoA in metazoan cells localizes to the cell cortex (Michaelson et al., 2001). Membrane-localization through the C-terminal CAAX motif is thought to increase Rho1 activity by increasing the likelihood of activation by GEFs. Therefore, we expected that the pool of Rho1-GTP we detect with the plasma-membrane localized Pkc1(HR1-C2)-mEGFP biosensor represents the relevant pool of active Rho1 in fission yeast.

The fluorescence of this probe is the same when free or bound to Rho1-GTP, so it can measure the local accumulation but not the total active Rho1-GTP. Binding of biosensors can compromise functions of the target protein, so expression of the biosensor may perturb cytokinesis if Rho1-GTP regulates furrow formation. To limit perturbation, the thiamine-repressible *3nmt1* promoter controlled the expression of Pkc1(HR1-C2)-mEGFP and cells were grown medium lacking thiamine (i.e. inducing conditions) for only ~15 h prior to imaging, the minimum time for maximum expression (Maundrell, 1990). Conveniently, expression from the *P3nmt1* promoter was highly variable, giving cells with a range of concentrations of Pkc1(HR1-C2)-mEGFP.

Expression of the biosensor consistently decreased the furrow ingression rate of wild-type, *mto2*_*S338N*_, and *mto1Δ* cells (Fig. 3F), delaying completion of constriction for all genotypes. On the other hand, the constriction rates were not strongly correlated with Pkc1(HR1-C2) expression level (by Spearman correlation test, which was used because Pkc1(HR1-C2) expression levels were not normally distributed)(Fig. S2B). The cytokinetic timelines of wild-type and *mto2*_*S338N*_ cells were statistically indistinguishable with and without Pkc1(HR1-C2), but expression of the biosensor delayed further both the duration of maturation and completion of constriction in *mto1Δ* cells (p < 0.05 by K-S test with Bonferroni correction) (Fig. 3D, E).

Fluorescence microscopy of Pkc1(HR1-C2)-mEGFP confirmed that Rho1-GTP concentrates at growing cell tips during interphase and at the cytokinetic furrow (Fig. 4A, Movie S1, Fig. 4A & B) as reported (Davidson et al., 2015). Pkc1(HR1-C2)-mEGFP also concentrated at other cortical locations; this ectopic localization was much more frequent in *mto2*_*S338N*_ (49%) and *mto1Δ* (46%) mutants than in wild-type (23%) cells (Fig. 4C, Movies S2, S3.). We only observed ectopic Pck1(HR1-C2) localization in the large fraction of cells lacking localization at cell ends (Fig. 4D). The *bgs1*_*D277N*_ cells expressing Pkc1(HR1-C2)-mEGFP grew very poorly at 36°C, so we were unable to quantify Rho1 activity. This genetic interaction suggests that *cps1-191* cells are more sensitive to the effects of the biosensor on Rho1 function. At 36°C a large fraction of cells of all strains did not concentrate the Pkc1(HR1-C2)-mEGFP biosensor at either cell end (nonpolar) or only accumulated the biosensor at one end (monopolar) (Fig. 4D). This phenotype was most severe in the *mto1Δ* mutant, which had the fewest bipolar cells and more nonpolar cells than in the wild type. This further indicates that expression of Pkc1(HR1-C2)- mEGFP compromises some cellular functions.

**Figure 4.**
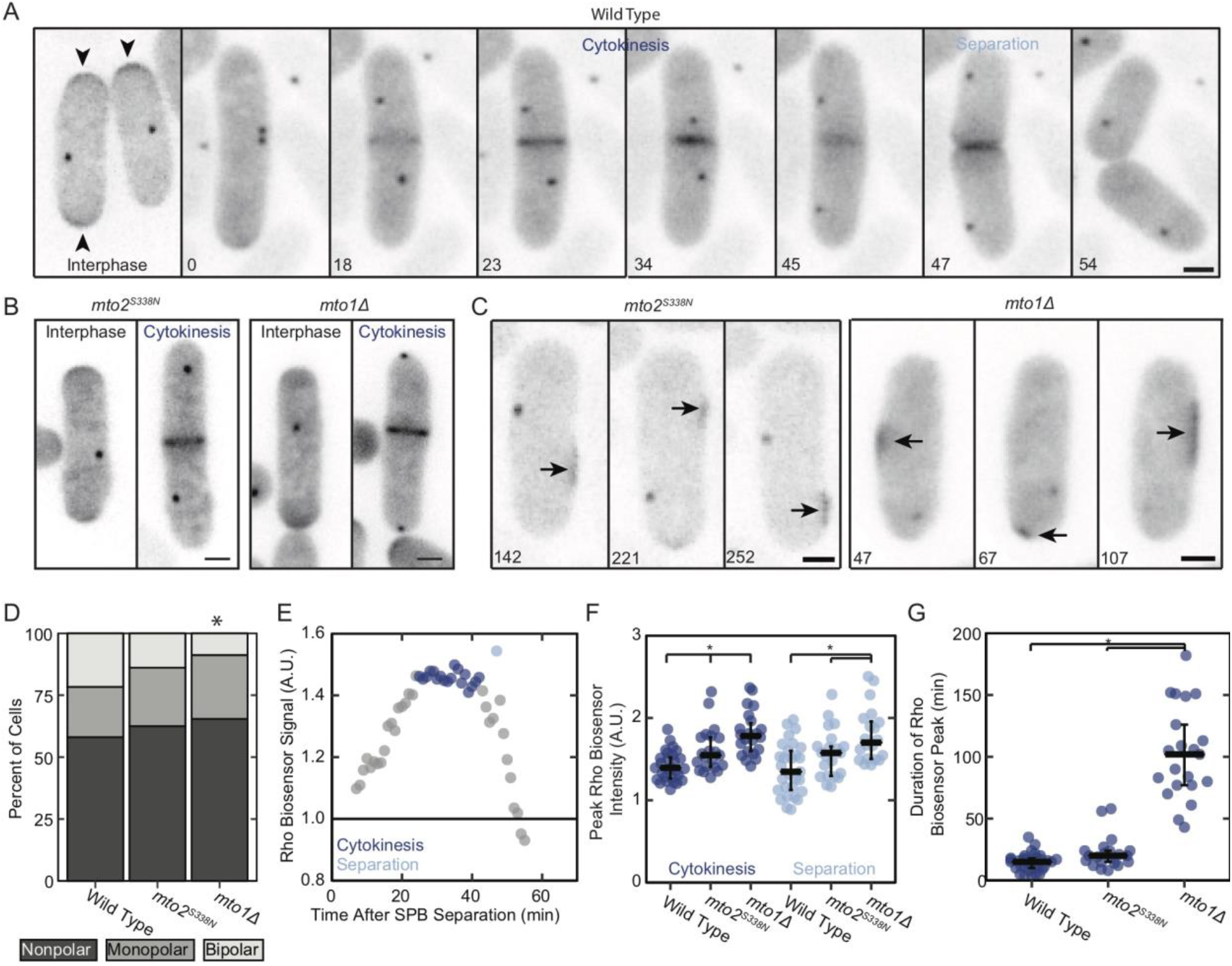
The *mto2*_*S338N*_ and *mto1Δ* mutations disrupt Rho1 signaling. A-C) Inverted-contrast maximum-intensity projected fluorescence micrographs of cells at 36°C expressing the Rho1 biosensor Pkc1(HR1-C2)-mEGFP and Pcp1-mEGFP to mark spindle pole bodies. Scale bars 2 μm. A) Wild-type cells. Active Rho1 concentrates at poles of interphase cells (arrowheads) and at the equator of cells during cytokinesis and septation. Times are minutes after SPB separation. B) *mto2*_*S338N*_ and *mto1Δ* cells exhibiting Rho1 concentration at cell ends during interphase (each left) and cell equators during cytokinesis (each right). C) Representative images of *mto1Δ* and *mto2*_*S338N*_ cells during interphase showing fluctuating, ectopic concentrations of the biosensor (arrows) at times denoted in minutes after the start of image acquisition. Ectopic accumulations were observed in 23% of wild-type, 49% of *mto2*_*S338N*_, and 46% of *mto1Δ* cells; n ≥ 72 cells, p = 0.007 and 0.006, respectively, from the wild-type population by Chi-Squared test with Bonferroni correction for multiple comparisons. D) The percent of cells exhibiting bipolar, monopolar, or nonpolar growth in an asynchronous population of cells expressing Pkc1(HR1-C2)-mEGFP shifted to 36°C. Asterisk denotes significant difference (p = 0.008) from the wild type as determined by Chi-Squared test with Bonferroni correction for multiple comparisons, n ≥ 143 cells. E) Time course of the integrated density of the Rho1 biosensor at equator of the cell in (A) during cytokinesis normalized to interphase value. The main peak (dark blue) is during furrow formation and the short second peak (light blue) is at cell separation. F & G) Rho1 biosensor signals at the equators of populations of ≥ 21 cells. Significance (p < 0.05) was determined using Welch’s ANOVA and Tukey’s post-hoc analysis. F) Peak Rho1 biosensor signal at the equator during cytokinesis and separation. G) Duration of Rho1 biosensor peak during cytokinesis.

The Pkc1(HR1-C2)-mEGFP signal peaked at the cleavage site during furrow ingression (Fig. 4C), followed by a brief decrease and second peak during cell separation (single light blue data point in Fig. 4C). The normalized peaks of Pkc1(HR1-C2)-mEGFP fluorescence during cytokinesis and at separation were more intense in *mto2*_*S338N*_ and *mto1Δ* cells than wild-type cells (Fig. 4D). Furthermore, the peak of Pkc1(HR1-C2)-mEGFP fluorescence during cytokinesis was prolonged in *mto1Δ* cells (Fig. 4E).

## Discussion

Since its description in 1999, the *cps1-191* strain has been the gold standard to generate *S. pombe* cells arrested with stable, unconstricting actomyosin rings. We find a more nuanced phenotype that depends on both the originally-identified *bgs1*_*D277N*_ mutation and an additional mutation, *mto2*_*S338N*_. Caution is advised when interpreting published results from experiments on *cps1-191* cells that examine the “arrest” phenotype, particularly those that required crossing the *cps1-191* strain with other strains, as the *mto2*_*S338N*_ mutation may no longer be present with the *bgs1*_*D277N*_ mutation. The *bgs1* and *mto2* loci are both on Chromosome II but are ~2 Mb apart. Isolating progeny that undergo a recombination event between these genes is common in our hands. A *cps1-191 mto2Δ* double mutant was previously examined, though the authors were presumably unaware of the *mto2*_*S338N*_ mutation in the *cps1-191* strain (Venkatram et al., 2005). However, we found that *bgs1*_*D277N*_ *mto2*_*S338N*_ had a more severe phenotype than *bgs1*_*D277N*_ *mto2Δ*. While the molecular basis of this difference is currently unknown, it may be that the *mto2*_*S338N*_ mutation exerts a dominant-negative effect on the function of Mto1 that is more severe than the entire loss of the Mto2 protein. We attempted to assess this hypothesis by ectopically introducing the *mto2*_*S338N*_ coding sequence in addition to the endogenous *mto2+* gene. However, progeny from crosses to generate a *mto2+*/*mto2*_*S338N*_ strain exhibited germination defects.

The *mto1Δ* and *mto2*_*S338N*_ cells recruit excess Rho1-GTP to the division site, but an additional insult (e.g. Bgs1 mutation, salt stress, Rho1 biosensor or GFP-Atb2 expression) is required to sensitize these cells and produce a furrow ingression phenotype. Why cells tolerate excess Rho1-GTP so well, and how increased Rho1-GTP modulates furrow ingression under multiple stressors are unclear. It might be expected that increased Rho1-GTP at the division sight would delay progression of cytokinesis, but *mto1Δ* and *mto2*_*S338N*_ cells exhibited wild-type cytokinesis timelines, which are normalized to SPB separation and mitotic timing. More work is required to assess which Rho1 regulators contribute to the observed hyperactivation and exactly how this influences furrow ingression. Many Rho1 regulators are known to be involved in septation, but how microtubules might regulate any of these proteins is currently unclear.

An additional mystery is why combination of Mto1/Mto2 mutations with the *bgs1*_*D227N*_ mutation decreases the furrowing rate while salt stress increases the furrowing rate, particularly because these mutants recruit normal levels of Bgs1 to the division site. Cells depleted of Bgs1 using the *nmt1* promoter and *cps1-191* cells exhibit abnormally-thick cell walls and secondary septa, suggesting that other glucan synthases attempt to compensate for the loss of Bgs1 activity (Cortés et al., 2007; 2015; Sethi et al., 2016). Excess Rho1-GTP at the division site may compromise the activity of these enzymes.

It was perplexing to us that an increase in recruitment of the Bgs1 regulator Rho1-GTP to the division site decreased the furrow ingression rate in *bgs1*_*D277N*_ cells. However, in mammalian cells, changing RhoA expression perturbs the levels of specific GTPases (e.g. Rac, Cdc42) through competition for shared GDP-dissociation inhibitors (GDIs), and it has been proposed that changes in the activity of RhoA could also affect additional GTPases through this mechanism (Boulter et al., 2010). *S. pombe* has a single Rho GDI, Gdi1, so an altered Rho1-GDP/GTP balance may change the amount of Gdi1 available to interact with other GTPases involved in cytokinesis such as Cdc42, Rho2, and Rho4. Rho1 activates Bgs1 (primary septum) and Bgs4 (secondary septum), but Rho2 regulates Ags1 (secondary septum). High Rho1 activity at the division site of *bgs1*_*D277N*_ *mto2*_*S338N*_ or *bgs1*_*D277N*_ *mto1Δ* cells may perturb Rho2 and Ags1 activity, compromising their ability to compensate for lack of Bgs1 and causing even slower furrow ingression than the *bgs1*_*D277N*_ substitution alone.

Our data suggest that Mto1 and Mto2 localized at the SPB participate in regulating furrow ingression. The SPB is known to serve as a hub for cell cycle signals—e.g. the SIN—and proteins localizing to the SPB throughout mitosis serve as markers for progression of the cell cycle (Tatebe et al., 2001; Grallert et al., 2004). Two, non-mutually-exclusive options are that Mto1/Mto2 and/or astral microtubules facilitate recruitment of known cell-cycle regulators to the SPB, or that astral microtubules may serve as signaling platforms in their own right. Cells lacking Mto1 have altered chromatid cohesion and DNA repair, likely due to loss of normal nuclear movement generated by microtubules (Zhurinsky, et al., 2019). These processes may be involved in the effects on furrow ingression; however, it is still unclear how nuclear status may be communicated to the division machinery independent of SIN. More research is required to determine exactly how microtubules influence the three systems required for furrow formation: an intact contractile ring; a signal originating from the cell cycle clock; and initiation of septum synthesis.

Our work demonstrates that crosstalk between microtubules and the furrow machinery mediated by Rho is conserved more broadly than previously thought and raises many intriguing questions regarding the molecular details of this communication in fungal cells. During cytokinesis of metazoan cells, RhoA regulates the location of the cytokinetic ring, activates formins for cytokinetic ring assembly, promotes myosin activity through Rho-associated protein kinase (ROCK), and promotes midbody ring maturation through Citron kinase (Pollard, 2017; El-Amine et al., 2019). Ring positioning is achieved through other mechanisms in fission yeast (Pollard, 2017), but Rho1 contributes to full activation of the SIN to drive ring formation (Alcaide-Gavilán et al., 2014) and has essential roles in septum synthesis through its interaction with glucan synthases (Arellano et al., 1996). This suggests that Rho signaling is an ancient mechanism for coordinating cytoskeletal elements and that during the divergence from the LECA (the last eukaryote common ancestor) Rho signaling has been adapted for a range of functions to regulate cytokinesis.

## Acknowledgments

Research reported in this publication was supported by National Institute of General Medical Sciences of the National Institutes of Health under award number R01GM026132 and an NRSA fellowship (F32GM125193) to S. Dundon. The content is solely the responsibility of the authors and does not necessarily represent the official views of the National Institutes of Health. The authors thank M. Balasubramanian, M. Das, and D. McCollum, for useful discussions; F. Chang, M. C. King, J. C. Ribas, K. E. Sawin, and J.-Q. Wu for generously sharing *S. pombe* strains; J. Berro and R. Fernandez for assistance with CRISPR; R. M. Dundon and T. J. Straub for assistance with genome analysis. The authors declare no competing financial interests.

## Methods and Materials

### Growth conditions and strain construction

Table S2 lists the *S. pombe* strains used in this work. Oligonucleotide primers were from Millipore Sigma. Restriction enzymes were from New England BioLabs. Plasmid purification was performed using an E.Z.N.A Plasmid Mini Kit (Omega Bio-tek) with Miraprep modifications (Pronobis et al., 2016). The Keck DNA Sequencing Lab of Yale University performed routine DNA sequencing. All cloning was done with high-efficiency NEB 5-alpha Competent *E. coli* from New England BioLabs. Standard methods were used for genetic crosses (Moreno et al., 1991).

*mto2*_*S338N*_:*KanMX6* and *mto2*_*S338C*_:*KanMX6* strains of *S. pombe* were generated by homologous integration (Bähler et al., 1998). First, pFA6a-mto2_S338N_-C-KanMX6 was generated. A 245-base fragment of the end of the *mto2* coding sequence was amplified from *cps1-191* (containing the *mto2*_*S338N*_ mutation) with flanking homology to the pFA6a vector using the ExpandTM High Fidelity PCR system (Millipore Sigma). The vector backbone was amplified from pFA6a-GFP-KanMX6 using Phusion polymerase (New England BioLabs). Both PCR reactions were digested with DpnI and purified using an E.Z.N.A Cycle Pure Kit (Omega Bio-tek). pFA6a-mto2_S338N_-C-KanMX6 was generated using NEBuilder HiFi DNA Assembly Master Mix (New England BioLabs) in an assembly containing 5 fmol of each fragment. The *mto2*_*S338N*_ segment was verified by sequencing. pFA6a-mto2_S338C_-C-KanMX6 was generated by site-directed mutagenesis. Mutagenic primers were used to amplify pFA6a-mto2_S338N_-C-KanMX6 using the Expand High Fidelity PCR system (Millipore Sigma). The PCR product was digested with DpnI and transformed into NEB 5-alpha Competent *E. coli*. The *mto2*_*S338C*_ segment was verified by sequencing.

PCR-based homologous integration cassettes were amplified using the ExpandTM High Fidelity PCR system (Millipore Sigma) from the appropriate plasmid with 80 bp flanking homology to replace the 3’ end of *mto2* (primers designed with PPPP at www.bahlerlab.info/resources/)(Bähler et al., 1998). Lithium acetate transformation was used to introduce 5-10 μg of integration cassette DNA (Murray et al., 2016). Integration at the correct locus was verified by colony PCR using LongAmp *Taq* 2X Master Mix (New England BioLabs), and the *mto2* gene was sequenced in its entirety to verify the S338N or S338C point mutations.

The *bgs1*_*D277N*_ strain was generated by CRISPR (Fernandez and Berro, 2016). A two-step PCR was used to assemble pUR19-adh1-Cas9-rrk1-sgRNA:Fex1+ with a guide sequence targeting *bgs1* using Q5 High-Fidelity polymerase (New England BioLabs). pUR19-adh1-Cas9-rrk1-sgRNA:Fex1+ and a 400-bp repair guide fragment containing the *bgs1*_*D277N*_ mutation were introduced into *fex1Δ fex2Δ S. pombe* cells by lithium acetate transformation (Murray et al., 2016). The *bgs1* gene was sequenced in its entirety to verify the *bgs1*_*D277N*_ mutation.

For serial dilution assays, 10 mL cultures of each strain were maintained in exponential phase for 36 hours in YE5S (rich) medium in 50 mL baffled flasks. All cultures were diluted to an OD_595_ of 0.5. Five 10-fold serial dilutions were made for each strain, yielding a total of six cellular densities. Five μL of each concentration was spotted on YE5S + 1.8% agar or YE5S 1.2 M Sorbitol + 1.8% agar plates. Each plate was grown either at 25°C or 36°C and imaged each day for four days to monitor growth.

### Whole-genome sequencing

*S. pombe* genomic DNA was isolated using a *Quick*-DNA Fungal/Bacterial Miniprep kit and purified by a Genomic DNA Clean & Concentrator-25 kit (both from Zymo Research). Genomic DNA was sequenced on a NovaSeq 6000 (Illumina) with 150 bp paired reads at the Yale Center for Genome Analysis. Sequence data quality was examined using FastQC v 0.11.7 (Babraham Bioinformatics). The *S. pombe* reference genome was downloaded from PomBase (Lock et al., 2019) and reads were aligned using MagicBLAST (Zhang et al., 2015). Aligned reads were sorted, duplicates removed, and indexed using SAMtools (Li et al., 2009). Deviations from the reference genome were identified using Pilon (Walker et al., 2014), and alignments were viewed using IGV 2.4.10 (Robinson et al., 2017). Identified mutations were cross-referenced with PomBase to determine location, eliminate previously-reported variants, and identify protein-coding mutations (Lock et al., 2019).

### Microscopy

Cells were grown in 50 mL baffled flasks in exponential phase at 25°C in YE5S (rich) medium for 24 h, then washed into EMM5S (low-fluorescence) medium for an additional 12 h. For experiments at 36°C, the culture was shifted to 36°C 2 h prior to mounting and imaging. To image, cells were isolated by centrifugation at 2300 × g for 30 s, washed with fresh EMM5S, and resuspended at ~20-fold higher concentration than in culture. Then 1.5-2.0 μL of concentrated cells were mounted on an EMM5S + 2% agarose pad freshly poured on a Therminator block (Davies et al., 2014) and a 24 × 50 mm no 1.5 coverslip was applied (Globe Scientific). Then 150 μL of EMM5S was injected beneath the coverslip in contact with the agarose pad and the edges of the chamber were sealed using VALAP to reduce shrinkage of the pad. For salt stress experiments, cells at 25°C were washed and resuspended in EMM5S + 0.6 M KCl, then immediately mounted on an EMM5S + 0.6 M KCl + 2% agarose pad on a Therminator block and imaged at 25°C. The average time between KCl addition and initiation of imaging was 5 minutes.

Cells were visualized using an Olympus IX-71 microscope with a 100×/numerical aperture (NA) 1.4 Plan Apo lens (Olympus) and a CSU-X1 (Andor Technology) confocal spinning-disk confocal system equipped with an iXON-EMCCD camera (Andor Technology). OBIS 488 nm LS 20 mW and OBIS 561 nm LS 20 mW lasers (Coherent) for excitation. The desired temperature was maintained during acquisition using a home-built version of the Therminator (Davies et al., 2014). Images were acquired using Micromanager 1.4 (Edelstein et al., 2014).

Acquisition settings for Pcp1-mEGFP Rlc1-tdTomato: 12 Z-slices were acquired with 0.5 μm spacing; 22 μW 561 nm laser power and 50 ms exposures; 35 μW 488 laser power and 50 ms exposures (power measured at the objective). An emission filter suitable for both tdTomato and mEGFP fluorescence was used to reduce the number of filter changes. All *bgs1+* cells were imaged at 4 XY positions and 1-min intervals. Cells carrying *bgs1*_*D277N*_ were imaged at 20 XY positions and 5-min intervals.

Acquisition settings for Rlc1-tdTomato Sfi1-mCherry mEGFP-Atb2: For cytokinesis timelines to determine the timing of spindle breakage, 12 Z-slices were acquired with 0.5 μm spacing at 3 XY positions and 1-minute intervals using 150 ms exposures with 32 μW 561 nm laser power and 200 ms exposures with 89 μW 488 laser power at the objective. Separate emission filters were used for tdTomato/mCherry and mEGFP fluorescence to avoid bleed through of the fluorophores with different spectral characteristics. To measure astral microtubule numbers and longevities, only mEGFP-Atb2 was imaged using 89 μW 488 laser power and 200 ms exposures of 10 Z-slices with 0.5 μm spacing at one XY position at 10-s intervals.

Acquisition settings for GFP-Bgs1 Rlc1-tdTomato Sfi1-mCherry: 12 Z-slices with 0.5 μm spacing were acquired at 3 XY positions and 1-min intervals using 150 ms exposures with 32 μW 561 nm laser power and 100 ms exposures with 89 μW 488 laser power at the objective. Separate emission filters were used for tdTomato/mCherry and mEGFP fluorescence to avoid bleed through of the fluorophores with different spectral characteristics. Strains bearing GFP-tagged proteins for the calibration curve were imaged under identical conditions. For camera noise correction, 100 images were acquired with 100 ms exposures and the shutter closed; for uneven illumination correction, dilute fluorescein was mounted on a 2% agarose EMM5S pad on the Therminator and 100 images were acquired using 50 ms exposures with 34 μW 488 laser power.

### Image Analysis

All image visualization and analyses were done using Fiji (Schindelin et al., 2012). Macros used for processing can be downloaded from https://github.com/SDundon/ImageJ-Processing-Macros. Prior to analysis, the contrast of each image was adjusted so that the black point was 5% below extracellular background signal and the white point was 20% above maximum signal intensity. Representative images were selected based on similarity to median values of each data set.

Constriction rates and cytokinesis timeline measurements: Maximum-intensity projections were produced for each time-lapse acquisition and a kymograph was first produced for each ring. If necessary, the StackReg plugin (Thévenaz et al., 1998) was used to register images across all time points. A line was drawn parallel to the ring, the “reslice” tool in Fiji used, and the resliced image was maximum-intensity projected to create a kymograph. Ring circumference for each time point was calculated using the 180912_AutoCircum.ijm macro. Circumference was plotted against time as a scatter plot in Microsoft Excel, and a linear trendline fit to the slope to calculate the constriction rate. For analysis of cells bearing the *bgs1*_*D277N*_ mutation (including *cps1-191*), cells were excluded from analysis if SPB separation occurred > 2 h after the start of acquisition. We found that rings formed after this time always constricted at an even slower rate, most likely due to effects of phototoxicity.

Cytokinesis milestones were defined as follows: “SPB separation” is the first frame in which two SPBs labeled with Pcp1-mEGFP could be resolved; “Complete assembly” is the first frame in which no Rlc1-tdTomato-labeled nodes were visible adjacent to the ring, because all had coalesced at the equator; “Constriction onset” is the first frame in which the Rlc1-tdTomato-labeled ring was smaller than its initial diameter and continued to decrease in size; “Constriction complete” is the frame in which the Rlc1-tdTomato signal reached its smallest size at the end of constriction. All milestones were temporally aligned with SPB separation set as time 0 and plotted as cumulative distributions. Spindle breakage was defined as the first time point in which there was no longer a single microtubule structure marked with GFP-Atb2 connecting the two SPBs. PAA appearance was defined as the first frame in which microtubules appeared at the equator distinct from the mitotic spindle.

Astral microtubule measurements: Astral microtubules were considered to be the same structure across consecutive frames (10 s intervals) if they a) originated from the same SPB and b) were oriented in the same direction ± 45° relative to the SPB in consecutive frames (Fig. 3A). The number of astral microtubules that originated from a single SPB during anaphase B was determined to exclude the intra-nuclear “astral” microtubules that form during earlier phases of mitosis (Zimmerman et al., 2004).

Rho biosensor and GFP-Bgs1 measurements: A ROI was selected for bleach correction reference using the following criteria: 1) must stay within the boundaries of one cell for the duration of the acquisition. 2) must be located within a cell that does not divide for the duration of the movie (to prevent differences in cytosolic concentration due to recruitment to the division site); 3) Pcp1-mEGFP-marked SPB must never enter the ROI. The same ROI was used for bleach correction of all channels. Bleach correction was conducted using the Fiji plugin for Exponential Fitting Method for the Bleach Correction. The intensity from all 12 Z-slices (0.5 μm apart) was then summed.

Rho biosensor measurements: Prior to analysis, the contrast of each image was adjusted so that the black point was 5% below extracellular background signal and the white point was 20% above maximum signal intensity of Pcp1-mEGFP, which was more consistent than pkc1(HR1-C2)-mEGFP signal. As previously observed, expression from the *3nmt1* promoter was highly variable. Therefore, cells were analyzed that had an average pkc1(HR1-C2)-mEGFP pixel intensity of >175,000 A.U. at SPB separation. This was calculated by drawing a spline-fit polygon ROI around the cell at the last time point prior to SPB separation. The integrated density was measured for this ROI, as well as a circular ROI covering the SPB (Fig. S2H). The SPB integrated density was subtracted from the cellular integrated density, and this value divided by the cell area with SPB area subtracted. This value (175,000 arbitrary units) was selected because cells of all genotypes below this threshold were found to have very low Peak Rho/Interphase Rho ratio values with a very different standard deviation than cells above this threshold. We reasoned that cells below this threshold express insufficient Pkc1(HR1-C2)-mEGFP to enable reliable measurement of Rho1-GTP enrichment at the cell equator. After elimination of cells below this threshold, the data were homoscedastic when the ratio was plotted against average pixel intensity.

To determine cellular polarity, a cell end was counted as ‘polarized’ if it exhibited visible Pkc1(HR1-C2)-mEGFP enrichment and/or was observed to grow over the course of the time-lapse experiment. Cells that exhibited no growth from either end were scored as ‘nonpolar,’ growth from one end was scored as ‘monopolar,’ and growth from both ends was ‘bipolar.’ Cells undergoing cytokinesis were excluded from this analysis, as cell ends cease growth during this time.

To calculate the Peak Rho/Interphase Rho ratio, a 3.75 × 1.95 μm ROI was positioned with the cytokinetic ring in the center (long axis to match the cell width)(Fig. S2H). The integrated density was measured for this ROI for each time point. These values were divided by the integrated density of this ROI for the time point immediately prior to SPB separation to normalize the intensity by Pkc1(HR1-C2)-mEGFP expression. Normalized intensity was plotted against time using Microsoft Excel, all time points in which the SPB was located within the ROI were excluded (Fig 3E). To measure peak duration, the peak fluorescence start was defined as the last time point of consistent signal increase across >3 time points. Peak end was defined as the last time point prior to consistent signal decrease below the peak start value across >3 time points. For cytokinesis peak signal, the maximum ratio between these points was determined. For separation peak signal, the maximum normalized signal within two time points of cell separation was determined.

Measurements of Bgs1 and Cdc7 molecule numbers: Bleach-corrected summed intensity images were corrected for camera noise and uneven illumination. To generate the calibration curve, a spline-fit polygon ROI was drawn around wild-type (unlabeled) cells and the average intensity in the ROI was measured (n = 74 cells). Subsequently, ROIs were similarly drawn around cells bearing seven different GFP-tagged proteins for the calibration curve (n > 65 cells). For each cell, the wild-type background fluorescence was subtracted and the integrated density was calculated. These integrated density measurements were plotted against the average number of molecules per cell for each protein reported in Wu and Pollard, 2005 and fit with a linear regression.

For Bgs1: A 1.0 μm by 3.75 μm (cell width) ROI was positioned to cover the equatorial GFP-Bgs1 signal and the integrated density measured from cell cycle time zero until 30 min past the completion of constriction. The same ROI was used to measure cytoplasmic GFP-Bgs1 signal adjacent to the ring at constriction onset. Cytoplasmic GFP-Bgs1 integrated density was subtracted from the equatorial signal until the onset of constriction. The average constriction rate and average ring size were used to calculate a correction factor for the progressively smaller cytoplasmic volume within the plane of the constricting ring, which was applied to the value of the cytoplasmic signal and subtracted from the equatorial signal until constriction completion.

For Cdc7: A 1.0 μm by 1.0 μm ROI was positioned over each SPB and the integrated density measured from the first time point where signal was visible above background (typically at SPB separation as determined by Sfi1-mCherry signal). The same ROI was used to measure cytoplasmic Cdc7-mEGFP at SPB separation, and this was subtracted from the signal measurements to correct for the large volume of cytoplasm not containing the SPB.

### Statistical analysis

All statistical tests were carried out in R (R Core Team, 2018). Plotting was done using tidyverse and ggbeeswarm packages (Wickham, 2017; Clarke and Sherrill-Mix, 2017), except for pie charts in Fig. 2, which were generated using Microsoft Excel.

## References

Alcaide-Gavilán, M., A. Lahoz, R.R. Daga, and J. Jimenez. 2014. Feedback regulation of SIN by Etd1 and Rho1 in fission yeast. Genetics. 196:455–470.

Arasada, R., and T.D. Pollard. 2014. Contractile ring stability in *S. pombe* depends on F-BAR protein Cdc15p and Bgs1p transport from the Golgi complex. Cell Rep. 8:1533–1544.

Arellano, M., A. Durán, and P. Pérez. 1996. Rho 1 GTPase activates the (1-3)β-D-glucan synthase and is involved in *Schizosaccharomyces pombe* morphogenesis. EMBO J. 15:4584–4591.

Balasubramanian, M.K., D. McCollum, L. Chang, K.C. Wong, N.I. Naqvi, X. He, S. Sazer, and K.L. Gould. 1998. Isolation and characterization of new fission yeast cytokinesis mutants. Genetics. 149:1265–1275.

Bähler, J., J.Q. Wu, M.S. Longtine, N.G. Shah, A. McKenzie, A.B. Steever, A. Wach, P. Philippsen, and J.R. Pringle. 1998. Heterologous modules for efficient and versatile PCR-based gene targeting in *Schizosaccharomyces pombe*. Yeast. 14:943–951.

Bezanilla, M., S.L. Forsburg, and T.D. Pollard. 1997. Identification of a second myosin-II in *Schizosaccharomyces pombe*: Myp2p is conditionally required for cytokinesis. Mol Biol Cell. 8:2693–2705.

Borek, W.E., L.M. Groocock, I. Samejima, J. Zou, F. de Lima Alves, J. Rappsilber, and K.E. Sawin. 2015. Mto2 multisite phosphorylation inactivates non-spindle microtubule nucleation complexes during mitosis. Nat Commun. 6:1–16.

Boulter, E., R. Garcia-Mata, C. Guilluy, A. Dubash, G. Rossi, P.J. Brennwald, and K. Burridge. 2010. Regulation of Rho GTPase crosstalk, degradation and activity by RhoGDI1. Nat Cell Biol. 12:477–483.

Calonge, T.M., K. Nakano, M. Arellano, R. Arai, S. Katayama, T. Toda, I. Mabuchi, and P. Pérez. 2000. *Schizosaccharomyces pombe* Rho2p GTPase regulates cell wall α-glucan biosynthesis through the protein kinase Pck2p. Mol Biol Cell. 11:4393–4401.

Cheffings, T.H., N.J. Burroughs, and M.K. Balasubramanian. 2019. Actin turnover ensures uniform tension distribution during cytokinetic actomyosin ring contraction. Mol Biol Cell. 30:933–941.

Chen, Q., and T.D. Pollard. 2011. Actin filament severing by cofilin is more important for assembly than constriction of the cytokinetic contractile ring. J Cell Biol. 195:485–498.

Clarke, E., and S. Sherrill-Mix. 2017. Categorical scatter (Violin point) plots [R package ggbeeswarm version 0.6.0].

Coffman, V.C., A.H. Nile, I.-J. Lee, H. Liu, and J.-Q. Wu. 2009. Roles of formin nodes and myosin motor activity in Mid1p-dependent contractile-ring assembly during fission yeast cytokinesis. Mol Biol Cell. 20:5195–5210.

Cortés, J.C.G., E. Carnero, J. Ishiguro, Y. Sánchez, A. Durán, and J.C. Ribas. 2005. The novel fission yeast (1,3)β-D-glucan synthase catalytic subunit Bgs4p is essential during both cytokinesis and polarized growth. J Cell Sci. 118:157–174.

Cortés, J.C.G., M. Konomi, I.M. Martins, J. Muñoz, M.B. Moreno, M. Osumi, A. Durán, and J.C. Ribas. 2007. The (1,3)β-d-glucan synthase subunit Bgs1p is responsible for the fission yeast primary septum formation. Mol Microbiol. 65:201–217.

Cortés, J.C.G., M. Sato, J. Muñoz, M.B. Moreno, J.A. Clemente-Ramos, M. Ramos, H. Okada, M. Osumi, A. Durán, and J.C. Ribas. 2012. Fission yeast Ags1 confers the essential septum strength needed for safe gradual cell abscission. J Cell Biol. 198:637–656.

Cortés, J.C.G., N. Pujol, M. Sato, M. Pinar, M. Ramos, B. Moreno, M. Osumi, J.C. Ribas, and P. Pérez. 2015. Cooperation between Paxillin-like protein Pxl1 and glucan synthase Bgs1 is essential for actomyosin ring stability and septum formation in fission yeast. PLoS Genet. 11:e1005358.

Davidson, R., D. Laporte, and J.-Q. Wu. 2015. Regulation of Rho-GEF Rgf3 by the arrestin Art1 in fission yeast cytokinesis. Mol Biol Cell. 26:453–466.

Davies, T., S.N. Jordan, V. Chand, J.A. Sees, K. Laband, A.X. Carvalho, M. Shirasu-Hiza, D.R. Kovar, J. Dumont, and J.C. Canman. 2014. High-resolution temporal analysis reveals a functional timeline for the molecular regulation of cytokinesis. Dev Cell. 30:209–223.

Dey, S.K., and T.D. Pollard. 2018. Involvement of the septation initiation network in events during cytokinesis in fission yeast. J Cell Sci. 131:jcs216895.

Edelstein, A.D., M.A. Tsuchida, N. Amodaj, H. Pinkard, R.D. Vale, and N. Stuurman. 2014. Advanced methods of microscope control using μManager software. J Biol Methods. 1:e10.

El-Amine, N., S.C. Carim, D. Wernike, and G.R.X. Hickson. 2019. Rho-dependent control of the Citron kinase, Sticky, drives midbody ring maturation. Mol Biol Cell. 30:2185–2204.

Fernandez, R., and J. Berro. 2016. Use of a fluoride channel as a new selection marker for fission yeast plasmids and application to fast genome editing with CRISPR/Cas9. Yeast. 33:549–557.

Grallert, A., A. Krapp, S. Bagley, V. Simanis, and I.M. Hagan. 2004. Recruitment of NIMA kinase shows that maturation of the *S. pombe* spindle-pole body occurs over consecutive cell cycles and reveals a role for NIMA in modulating SIN activity. Genes Dev. 18:1007–1021.

Janson, M.E., T.G. Setty, A. Paoletti, and P.T. Tran. 2005. Efficient formation of bipolar microtubule bundles requires microtubule-bound gamma-tubulin complexes. J Cell Biol. 169:297–308.

Kono, K., Y. Saeki, S. Yoshida, K. Tanaka, and D. Pellman. 2012. Proteasomal degradation resolves competition between cell polarization and cellular wound healing. Cell. 150:151–164.

Laplante, C., J. Berro, E. Karatekin, A. Hernandez-Leyva, R. Lee, and T.D. Pollard. 2015. Three myosins contribute uniquely to the assembly and constriction of the fission yeast cytokinetic contractile ring. Curr Biol. 25:1955–1965.

Leong, S.L., E.M. Lynch, J. Zou, Y.D. Tay, W.E. Borek, M.W. Tuijtel, J. Rappsilber, and K.E. Sawin. 2019. Reconstitution of microtubule nucleation *in vitro* reveals novel roles for Mzt1. Curr Biol. 29:2199–2207.e10.

Li, H., B. Handsaker, A. Wysoker, T. Fennell, J. Ruan, N. Homer, G. Marth, G. Abecasis, R. Durbin, 1000 Genome Project Data Processing Subgroup. 2009. The Sequence Alignment/Map format and SAMtools. Bioinformatics. 25:2078–2079.

Li, Y., J.R. Christensen, K.E. Homa, G.M. Hocky, A. Fok, J.A. Sees, G.A. Voth, and D.R. Kovar. 2016. The F-actin bundler α-actinin Ain1 is tailored for ring assembly and constriction during cytokinesis in fission yeast. Mol Biol Cell. 27:1821–1833.

Liu, J., H. Wang, D. McCollum, and M.K. Balasubramanian. 1999. Drc1p/Cps1p, a 1,3-β-glucan synthase subunit, is essential for division septum assembly in *Schizosaccharomyces pombe*. Genetics. 153:1193–1203.

Lock, A., K. Rutherford, M.A. Harris, J. Hayles, S.G. Oliver, J. Bahler, and V. Wood. 2019. PomBase 2018: user-driven reimplementation of the fission yeast database provides rapid and intuitive access to diverse, interconnected information. Nucleic Acids Res. 47:D821–D827.

Loo, T.-H., and M. Balasubramanian. 2008. *Schizosaccharomyces pombe* Pak-related protein, Pak1p/Orb2p, phosphorylates myosin regulatory light chain to inhibit cytokinesis. J Cell Biol. 183:785–793.

Maundrell, K. 1990. nmt1 of fission yeast. A highly transcribed gene completely repressed by thiamine. J. Biol. Chem. 265:10857–10864.

Michaelson, D., J. Silletti, G. Murphy, P. D’Eustachio, M. Rush, and M.R. Philips. 2001. Differential localization of Rho GTPases in live cells: regulation by hypervariable regions and RhoGDI binding. J Cell Biol. 152:111–126.

Minet, M., P. Nurse, P. Thuriaux, and J.M. Mitchison. 1979. Uncontrolled septation in a cell division cycle mutant of the fission yeast *Schizosaccharomyces pombe*. J Bacteriol. 137:440–446.

Mitchison, J.M., and P. Nurse. 1985. Growth in cell length in the fission yeast *Schizosaccharomyces pombe*. J Cell Sci. 75:357–376.

Morris, Z., D. Sinha, A. Poddar, B. Morris, and Q. Chen. 2019. Fission yeast TRP channel Pkd2p localizes to the cleavage furrow and regulates cell separation during cytokinesis. Mol Biol Cell. 30:1791–1804.

Moreno, S., A. Klar, and P. Nurse. 1991. Molecular genetic analysis of fission yeast Schizosaccharomyces pombe. Meth. Enzymol. 194:795–823.

Murray, J.M., A.T. Watson, and A.M. Carr. 2016. Transformation of *Schizosaccharomyces pombe*: lithium acetate/dimethyl sulfoxide procedure. Cold Spring Harbor Protocols. 2016:pdb.prot090969.

Nakano, K., R. Arai, and I. Mabuchi. 1997. The small GTP-binding protein Rho1 is a multifunctional protein that regulates actin localization, cell polarity, and septum formation in the fission yeast *Schizosaccharomyces pombe*. Genes Cells. 2:679–694.

Okada, H., C. Wloka, J.-Q. Wu, and E. Bi. 2019. Distinct roles of Myosin-II isoforms in cytokinesis under normal and stressed conditions. ISCIENCE. 14:69–87.

Pardo, M., and P. Nurse. 2003. Equatorial retention of the contractile actin ring by microtubules during cytokinesis. Science. 300:1569–1574.

Pelham, R.J., and F. Chang. 2002. Actin dynamics in the contractile ring during cytokinesis in fission yeast. Nature. 419:82–86.

Pollard, T.D. 2017. Nine unanswered questions about cytokinesis. J Cell Biol. 216:3007–3016.

Proctor, S.A., N. Minc, A. Boudaoud, and F. Chang. 2012. Contributions of turgor pressure, the contractile ring, and septum assembly to forces in cytokinesis in fission yeast. Curr Biol. 22:1601–1608.

Pronobis, M.I., N. Deuitch, and M. Peifer. 2016. The Miraprep: A protocol that uses a Miniprep kit and provides Maxiprep yields. PLoS ONE. 11:e0160509.

Roberts-Galbraith, R.H., J.-S. Chen, J. Wang, and K.L. Gould. 2009. The SH3 domains of two PCH family members cooperate in assembly of the *Schizosaccharomyces pombe* contractile ring. J Cell Biol. 184:113–127.

Roberts-Galbraith, R.H., M.D. Ohi, B.A. Ballif, J.-S. Chen, I. McLeod, W.H. McDonald, S.P. Gygi, J.R. Yates, and K.L. Gould. 2010. Dephosphorylation of F-BAR protein Cdc15 modulates its conformation and stimulates its scaffolding activity at the cell division site. Mol Cell. 39:86–99.

Robinson, J.T., H. Thorvaldsdóttir, A.M. Wenger, A. Zehir, and J.P. Mesirov. 2017. Variant review with the Integrative Genomics Viewer. Cancer Res. 77:e31–e34.

Samejima, I., P.C.C. Lourenco, H.A. Snaith, and K.E. Sawin. 2005. Fission yeast Mto2p regulates microtubule nucleation by the centrosomin-related protein Mto1p. Mol Biol Cell. 16:3040–3051.

Samejima, I., V.J. Miller, S.A. Rincón, and K.E. Sawin. 2010. Fission yeast Mto1 regulates diversity of cytoplasmic microtubule organizing centers. Curr Biol. 20:1959–1965.

Sawin, K.E., and P.T. Tran. 2006. Cytoplasmic microtubule organization in fission yeast. Yeast. 23:1001–1014.

Sawin, K.E., P.C.C. Lourenco, and H.A. Snaith. 2004. Microtubule nucleation at non-spindle pole body microtubule-organizing centers requires fission yeast centrosomin-related protein mod20p. Curr Biol. 14:763–775.

Schindelin, J., I. Arganda-Carreras, E. Frise, V. Kaynig, M. Longair, T. Pietzsch, S. Preibisch, C. Rueden, S. Saalfeld, B. Schmid, J.-Y. Tinevez, D.J. White, V. Hartenstein, K. Eliceiri, P. Tomancak, and A. Cardona. 2012. Fiji: an open-source platform for biological-image analysis. Nat Meth. 9:676–682.

Sethi, K., S. Palani, J.C.G. Cortés, M. Sato, M. Sevugan, M. Ramos, S. Vijaykumar, M. Osumi, N.I. Naqvi, J.C. Ribas, and M. Balasubramanian. 2016. A new membrane protein Sbg1 links the contractile ring apparatus and septum synthesis machinery in fission yeast. PLoS Genet. 12:e1006383–30.

Simanis, V. 2015. Pombe’s thirteen - control of fission yeast cell division by the septation initiation network. J Cell Sci. 128:1465–1474.

Sohrmann, M., S. Schmidt, I. Hagan, and V. Simanis. 1998. Asymmetric segregation on spindle poles of the *Schizosaccharomyces pombe* septum-inducing protein kinase Cdc7p. Genes Dev. 12:84–94.

Tatebe, H., G. Goshima, K. Takeda, T. Nakagawa, K. Kinoshita, and M. Yanagida. 2001. Fission yeast living mitosis visualized by GFP-tagged gene products. Micron. 32:67–74.

R Core Team. R: A language and environment for statistical computing. 2018. R Foundation for Statistical Computing.

Tebbs, I.R., and T.D. Pollard. 2013. Separate roles of IQGAP Rng2p in forming and constricting the *Schizosaccharomyces pombe* cytokinetic contractile ring. Mol Biol Cell. 24:1904–1917.

Thévenaz, P., U.E. Ruttimann, and M. Unser. 1998. A pyramid approach to subpixel registration based on intensity. IEEE Trans Image Process. 7:27–41.

Thiyagarajan, S., E.L. Munteanu, R. Arasada, T.D. Pollard, and B. O’Shaughnessy. 2015. The fission yeast cytokinetic contractile ring regulates septum shape and closure. J Cell Sci. 128:3672–3681.

Vavylonis, D., J.-Q. Wu, S. Hao, B. O’Shaughnessy, and T.D. Pollard. 2008. Assembly mechanism of the contractile ring for cytokinesis by fission yeast. Science. 319:97–100.

Venkatram, S., J.L. Jennings, A. Link, and K.L. Gould. 2005. Mto2p, a novel fission yeast protein required for cytoplasmic microtubule organization and anchoring of the cytokinetic actin ring. Mol Biol Cell. 16:3052–3063.

Wachtler, V., Y. Huang, J. Karagiannis, and M.K. Balasubramanian. 2006. Cell cycle-dependent roles for the FCH-domain protein Cdc15p in formation of the actomyosin ring in *Schizosaccharomyces pombe*. Mol Biol Cell. 17:3254–3266.

Walker, B.J., T. Abeel, T. Shea, M. Priest, A. Abouelliel, S. Sakthikumar, C.A. Cuomo, Q. Zeng, J. Wortman, S.K. Young, and A.M. Earl. 2014. Pilon: an integrated tool for comprehensive microbial variant detection and genome assembly improvement. PLoS ONE. 9:e112963.

Wang, N., L. Lo Presti, Y.-H. Zhu, M. Kang, Z. Wu, S.G. Martin, and J.-Q. Wu. 2014. The novel proteins Rng8 and Rng9 regulate the myosin-V Myo51 during fission yeast cytokinesis. J Cell Biol. 205:357–375.

Wickham, H. 2017. tidyverse: Easily Install and Load the “Tidyverse.”

Wood, V., R. Gwilliam, M.-A. Rajandream, M. Lyne, R. Lyne, A. Stewart, J. Sgouros, N. Peat, J. Hayles, S. Baker, D. Basham, S. Bowman, K. Brooks, D. Brown, S. Brown, T. Chillingworth, C. Churcher, M. Collins, R. Connor, A. Cronin, P. Davis, T. Feltwell, A. Fraser, S. Gentles, A. Goble, N. Hamlin, D. Harris, J. Hidalgo, G. Hodgson, S. Holroyd, T. Hornsby, S. Howarth, E.J. Huckle, S. Hunt, K. Jagels, K. James, L. Jones, M. Jones, S. Leather, S. McDonald, J. McLean, P. Mooney, S. Moule, K. Mungall, L. Murphy, D. Niblett, C. Odell, K. Oliver, S. O’Neil, D. Pearson, M.A. Quail, E. Rabbinowitsch, K. Rutherford, S. Rutter, D. Saunders, K. Seeger, S. Sharp, J. Skelton, M. Simmonds, R. Squares, S. Squares, K. Stevens, K. Taylor, R.G. Taylor, A. Tivey, S. Walsh, T. Warren, S. Whitehead, J. Woodward, G. Volckaert, R. Aert, J. Robben, B. Grymonprez, I. Weltjens, E. Vanstreels, M. Rieger, M. Schäfer, S. Müller-Auer, C. Gabel, M. Fuchs, A. Düsterhöft, C. Fritzc, E. Holzer, D. Moestl, H. Hilbert, K. Borzym, I. Langer, A. Beck, H. Lehrach, R. Reinhardt, T.M. Pohl, P. Eger, W. Zimmermann, H. Wedler, R. Wambutt, B. Purnelle, A. Goffeau, E. Cadieu, et al. 2002. The genome sequence of *Schizosaccharomyces pombe*. Nature. 415:871–880.

Wu, J.-Q., J.R. Kuhn, D.R. Kovar, and T.D. Pollard. 2003. Spatial and temporal pathway for assembly and constriction of the contractile ring in fission yeast cytokinesis. Dev Cell. 5:723–734.

Yamashita, A., M. Sato, A. Fujita, M. Yamamoto, and T. Toda. 2005. The roles of fission yeast ase1 in mitotic cell division, meiotic nuclear oscillation, and cytokinesis checkpoint signaling. Mol Biol Cell. 16:1378–1395.

Zhang, W., Y. Yu, F. Hertwig, J. Thierry-Mieg, W. Zhang, D. Thierry-Mieg, J. Wang, C. Furlanello, V. Devanarayan, J. Cheng, Y. Deng, B. Hero, H. Hong, M. Jia, L. Li, S.M. Lin, Y. Nikolsky, A. Oberthuer, T. Qing, Z. Su, R. Volland, C. Wang, M.D. Wang, J. Ai, D. Albanese, S. Asgharzadeh, S. Avigad, W. Bao, M. Bessarabova, M.H. Brilliant, B. Brors, M. Chierici, T.-M. Chu, J. Zhang, R.G. Grundy, M.M. He, S. Hebbring, H.L. Kaufman, S. Lababidi, L.J. Lancashire, Y. Li, X.X. Lu, H. Luo, X. Ma, B. Ning, R. Noguera, M. Peifer, J.H. Phan, F. Roels, C. Rosswog, S. Shao, J. Shen, J. Theissen, G.P. Tonini, J. Vandesompele, P.-Y. Wu, W. Xiao, J. Xu, W. Xu, J. Xuan, Y. Yang, Z. Ye, Z. Dong, K.K. Zhang, Y. Yin, C. Zhao, Y. Zheng, R.D. Wolfinger, T. Shi, L.H. Malkas, F. Berthold, J. Wang, W. Tong, L. Shi, Z. Peng, and M. Fischer. 2015. Comparison of RNA-seq and microarray-based models for clinical endpoint prediction. Genome Biol. 16:133.

Zhou, Z., E.L. Munteanu, J. He, T. Ursell, M. Bathe, K.C. Huang, and F. Chang. 2015. The contractile ring coordinates curvature-dependent septum assembly during fission yeast cytokinesis. Mol Biol Cell. 26:78–90.

Zhurinsky, J., S. Salas-Pino, A.B. Iglesias-Romero, A. Torres-Mendez, B. Knapp, I. Flor-Parra, J. Wang, K. Bao, S. Jia, F. Chang, and R.R. Daga. (2019). Effects of the microtubule nucleator Mto1 on chromosomal movement, DNA repair, and sister chromatid cohesion in fission yeast. Mol Biol Cell 30, 2695–2708.

Zimmerman, S., and F. Chang. 2005. Effects of γ-tubulin complex proteins on microtubule nucleation and catastrophe in fission yeast. Mol Biol Cell. 16:2719–2733.

Zimmerman, S., R.R. Daga, and F. Chang. 2004. Intra-nuclear microtubules and a mitotic spindle orientation checkpoint. Nat Cell Biol. 6:1245–1246.

