## Supplemental Materials for "Microtubule Nucleation Promoters Mto1 and Mto2 Regulate Cytokinesis in Fission Yeast"

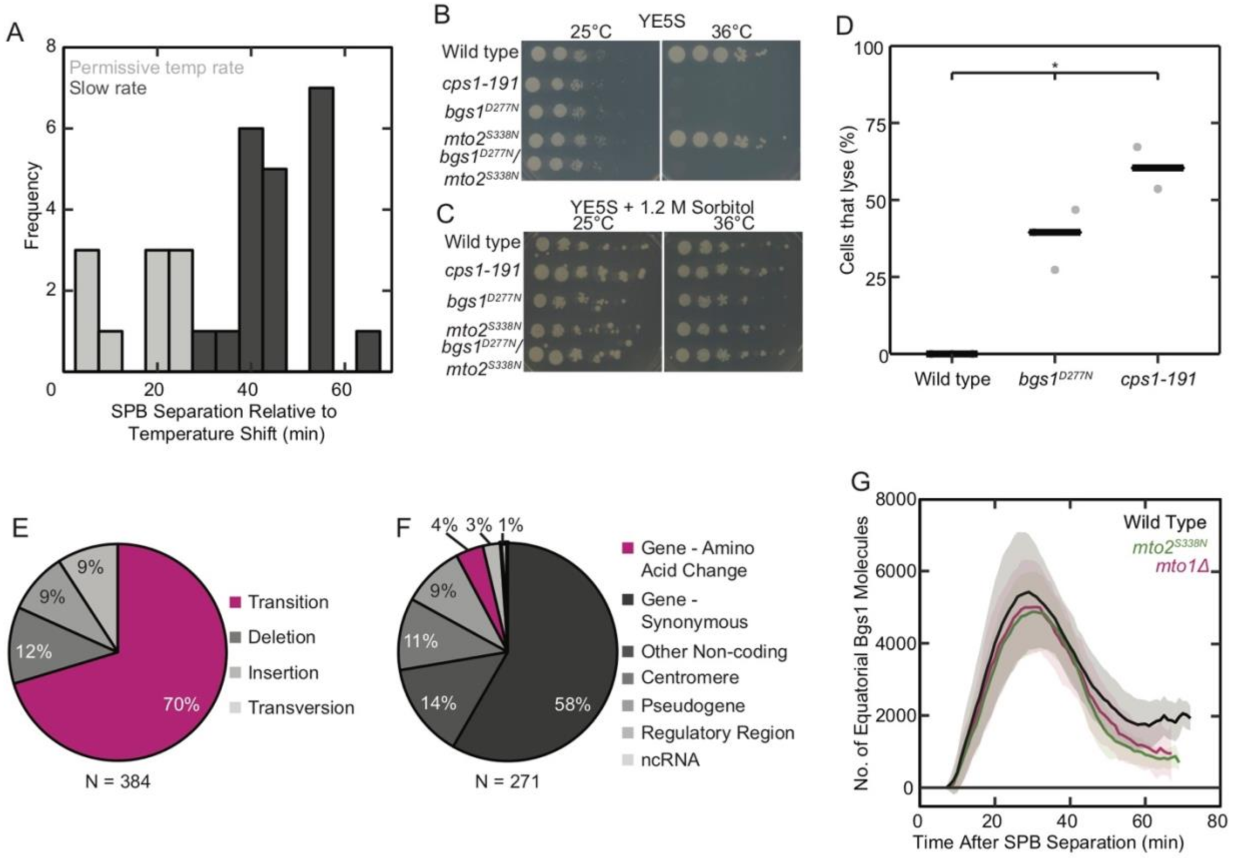


**Figure S1. The *cps1-191* constriction phenotype arises after ~30 minutes, this strain contains a large number of mutations, and cells die by lysis. Neither *mto2*$_{S338N}$ nor *mto1Δ* cells have defects in recruiting Bgs1 to the equator.** A) Histogram of *cps1-191* cells that constrict normally or at a slow rate after shift from permissive (25°C) to restrictive (36°C) temperature, n = 31 cells. B & C) Serial-dilution growth assays comparing the temperature sensitivity of strains carrying mutations of *bgs1*$_{+}$ and *mto2*$_{+}$. B) YE5S agar plate. C) YE5S + 1.2 M sorbitol agar plate. D) Lysis of wild-type, *bgs1*$_{D277N}$, and *cps1-191* strains. Each point is the average of n ≥ 13 cells that lysed during one multi-point image acquisition. E) Categorization of mutation types in *cps1-191*. F) Locations of *cps1-191* transition mutations. G) Time course of the accumulation of mGFP-Bgs1 molecules at the cell equator during cytokinesis. Averages of samples of 6 or more cells. The peak values were not significantly different by Welch's ANOVA.

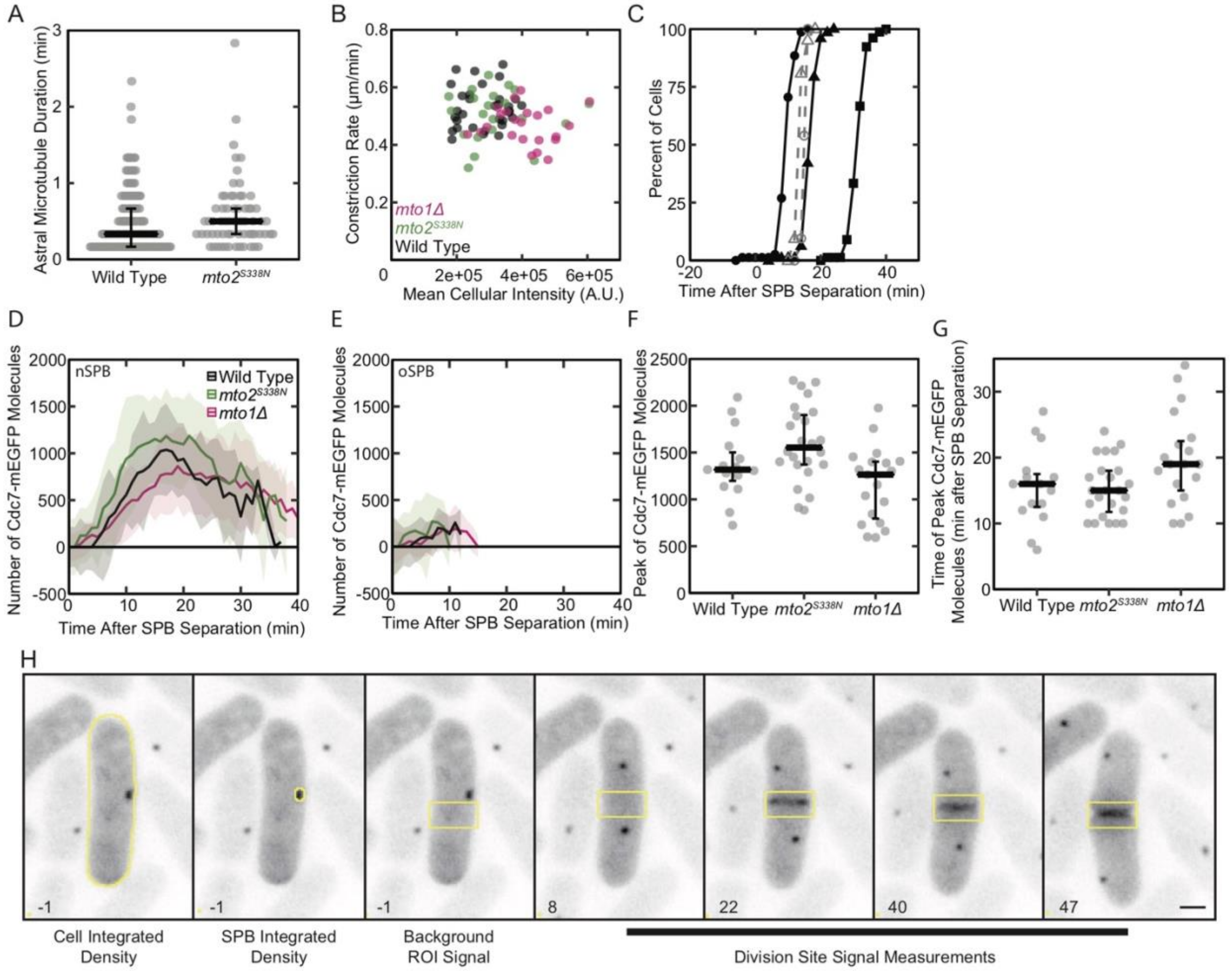


**Figure S2. Astral microtubules in *mto2*$_{S338N}$ cells last as long as in wild-type cells, Pkc1(H1-C2)-mEGFP expression does not correlate with constriction rate, and timing of spindle breakage and PAA formation relative to the cytokinetic timeline in wild-type cells.** A) Durations of single astral microtubules attached to a SPB during anaphase B, n ≥ 68 microtubules. The median and 1st and 3rd quartiles are indicated by black bars. Samples were not significantly different based on a K-S test. B) Dependence of the cytokinetic ring constriction rate on the mean fluorescence intensity of Pkc1(HR1-C2)-mEGFP of n ≥ 21 cells per strain. C) Comparison of contractile ring milestones (black, solid lines) with mitotic microtubule events (gray, dashed lines) in samples of ≥ 21 wild-type cells at 36°C. Cumulative distribution plot of cells with (●) assembled contractile rings, (■) constricting contractile rings, (▲) constricted rings, (○) broken spindles, and (△) formed PAA. D & E) Time course of the recruitment of Cdc7-mEGFP molecules to the new (D) and old (E) SPB. F) The peak number of Cdc7-mEGFP molecules at the nSPB. G) The time after SPB separation when the number of Cdc7-mEGFP molecules peaked. For G & H, neither mutant strain differed from the wild-type values by Welch's ANOVA, $n \geq 15$ SPBs for E-G. H) Inverted-contrast maximum-intensity projected fluorescence micrographs of a representative cell at 36°C expressing the Rho1 biosensor Pkc1(HR1-C2)-mEGFP and Pcp1-mEGFP to mark spindle pole bodies. Scale bar 2 μm, time annotated relative to SPB separation, ROIs depicted in yellow. To quantify Rho1-GTP recruitment each image was bleach corrected, sum projected, and quantitatively contrasted. The integrated density of the entire cell was measured at t = -1, and the integrated density of the SPB at the same time point was subtracted from this value to calculate the average signal per pixel. This was used to determine whether Pkc1(HR1-C2)-mEGFP signal was sufficiently high for valid measurements. A 3.75 x 1.95 μm ROI was positioned as close to the cell center as possible while excluding the SPB at t = -1 and the integrated density measured. The same ROI was then positioned over the division site and the integrated density was measured at each time point through cytokinesis. Each value was divided by the integrated density measured at t = -1 to normalize for the variable Pkc1(HR1-C2)-mEGFP expression levels. These values were plotted over time as in Fig. 3E.

**Supplemental movies**

**Movie S1.** Rho biosensor Pkc1(HR1-C2)-mEGFP localization during wild-type *S. pombe* cytokinesis. Time expressed relative to SPB separation, scale 2 μm.

**Movie S2.** Rho biosensor Pkc1(HR1-C2)-mEGFP localization in interphase *mto2*$_{S338N}$ mutant cells. Time expressed relative to acquisition start, scale 2 μm.

**Movie S3.** Rho biosensor Pkc1(HR1-C2)-mEGFP localization in interphase *mto1Δ* mutant cells. Time expressed relative to acquisition start, scale 2 μm.

**Table S1. Correlation between Mto2 and Nup40 mutations with constriction rate phenotype.** Progeny from *cps1-191* crosses that were confirmed to contain the *bgs1*$_{D277N}$ mutation were examined to determine whether they exhibited a furrow ingression rate similar to *cps1-191* or *bgs1*$_{D277N}$ cells ("Phenotype" column). The *mto2* and *nup40* loci were sequenced to determine whether each strain carried a wild-type version of the gene or the mutant version identified in the *cps1-191* strain. The slowest ingression rate (*cps1-191* phenotype) specifically correlated with the presence of the *mto2*$_{S338N}$ mutation, and did not correlate with the *nup40*$_{T281I}$ mutation.

| Strain ID | Mto2 | Nup40 | Phenotype |
|---|---|---|---|
| SDP 014 | S338N | T281I | *cps1-191* |
| SDP 062.1 | S338N | Wild type | *cps1-191* |
| SDP 062.2 | S338N | Wild type | *cps1-191* |
| SDP 162.3 | S338N | Wild type | *cps1-191* |
| SDP 162.5 | S338N | Wild type | *cps1-191* |
| SDP 162.6 | S338N | Wild type | *cps1-191* |
| SDP 062.3 | Wild type | T281I | *bgs1*$^{D277N}$ |
| SDP 162.1 | Wild type | Wild type | *bgs1*$^{D277N}$ |
| SDP 162.2 | Wild type | Wild type | *bgs1*$^{D277N}$ |

**Table S2. *S. pombe* strains used in this study.**

| Strain ID | Genotype | Source |
|---|---|---|
| SDP 014 | *h- cps1-191 leu1-32 ura4-Δ18* | JR 1024, Liu et al 1999 |
| SDP 016 | *h- ade6-M210 leu1-32 ura4-Δ18* | Pollard lab stock |
| SDP 054.1 | *h- Rlc1-tdTomato:NatMX6 Pcp1-GFP:KanR ura4-Δ18 leu1-32 ade6-M210* | This study |
| SDP 062.1 | *h- cps1-191 Rlc1-tdTomato:NatMX6 Pcp1-GFP:KanMX6 leu1-32 ura4-Δ18 ade6-M21x* | This study |
| SDP 062.2 | *h- cps1-191 Rlc1-tdTomato:NatMX6 Pcp1-GFP:KanR leu1-32 ura4-Δ18 ade6-M21x* | This study |
| SDP 062.3 | *h- cps1-191 Rlc1-tdTomato:NatMX6 Pcp1-GFP:KanR leu1-32 ura4-Δ18 ade6-M210* | This study |
| SDP 131 | *h- arc5-mGFP:KanMX6 leu1-32 ura4-D18 his3-D1 ade6-M216* | Wu and Pollard 2005 |
| SDP 134 | *h+ fim1-mGFP:kanMX6 leu1-32 ura4-D18 ade6-M210* | Wu and Pollard 2005 |
| SDP 142 | *h+ acp2-mGFP:KanMX6 ade6-M216 his3-Δ1 leu1-32 ura4-Δ18* | Wu and Pollard 2005 |
| SDP 143 | *h+ ain1-mGFP:KanMX6 ade6-M216 his3-Δ1 leu1-32 ura4-Δ18* | Wu and Pollard 2005 |
| SDP 144 | *h+ arp2-mGFP:KanMX6 ade6-M216 his3-Δ1 leu1-32 ura4-Δ18* | Wu and Pollard 2005 |
| SDP 145 | *h+ arp3-mGFP:KanMX6 ade6-M216 his3-Δ1 leu1-32 ura4-Δ18* | Wu and Pollard 2005 |
| SDP 146 | *h+ KanMX6:Pmyo2-mGFP-myo2 ade6-M216 his3-Δ1 leu1-32 ura4-Δ18* | Wu and Pollard 2005 |
| SDP 162.1 | *h+ cps1-191 Rlc1-tdTomato:NatMX6 Pcp1-GFP:KanMX6 ura4-Δ18 ade6-M21x* | This study |
| SDP 162.2 | *h+ cps1-191 Rlc1-tdTomato:NatMX6 Pcp1-GFP:KanMX6 ura4-Δ18 ade6-M21x* | This study |
| SDP 162.3 | *h+ cps1-191 Rlc1-tdTomato:NatMX6 Pcp1-GFP:KanMX6 ura4-Δ18 ade6-M21x* | This study |
| SDP 162.5 | *h+ cps1-191 Rlc1-tdTomato:NatMX6 Pcp1-GFP:KanMX6 ura4-Δ18 ade6-M21x* | This study |
| SDP 162.6 | *h+ cps1-191 Rlc1-tdTomato:NatMX6 Pcp1-GFP:KanMX6 ura4-Δ18 ade6-M21x* | This study |
| SDP 224.1 | *h+ mto2S338N:KanMX6 ade6-M216 his3-Δ1 leu1-32 ura4-Δ18* | This study |
| SDP 241 | *h+ mto1Δ::kanMX6 ade6-216 leu1-32 ura4-Δ18* | KS 1017, Lynch, et al. 2014 |
| SDP 243 | *h+ mto2Δ::kanMX6 ade6-M216 leu1-32 ura4-Δ18* | KS 977, Samejima, et al. 2005 |
| SDP 247 | *h+ mto2S338N:KanMX6 Rlc1-tdTomato:NatMX6 Pcp1-mEGFP:KanMX6 ura4-Δ18 leu1-32 ade6-M21x* | This study |
| SDP 248 | *h- mto2Δ:KanMX6 Rlc1-tdTomato:NatMX6 Pcp1-mEGFP:KanMX6 ade6-M21x leu1-32 ura4-Δ18* | This study |
| SDP 250.1 | *h- mto1Δ:KanMX6 Rlc1-tdTomato:NatMX6 Pcp1-mEGFP:KanMX6 ade6-M21x leu1-32 ura4-Δ18* | This study |
| SDP 254.1 | *h- bgs1D277N fex1Δ fex2Δ ade6-M216 his3-Δ1 leu1-32 ura4-Δ18* | This study |
| SDP 258 | *h- mto2S338N:KanMX6 bgs1D277N Rlc1-tdTomato:NatMX6 Pcp1-mEGFP:KanMX6 ade6-M21x his3-Δ1 leu1-32 ura4-Δ18 fex1? fex2?* | This study |

| Strain ID | Genotype | Source |
|---|---|---|
| SDP 259.1 | *h- bgs1D277N Rlc1-tdTomato:NatMX6 Pcp1-GFP:KanMX6 ade6-M21x leu1-32 ura4-Δ18 fex1? fex2?* | This study |
| SDP 266 | *h+ mto2S338C:KanMX6 Rlc1-tdTomato:NatMX6 Pcp1-mEGFP:KanMX6 ade6-M21x his3-Δ1 leu1-32 ura4-Δ18* | This study |
| SDP 328.1 | *h- mto2Δ::KanMX6 bgs1D277N Rlc1-tdTomato:NatMX6 Pcp1-GFP:KanMX6 ade6-M21x his3-Δ1 leu1-32 ura4-Δ18 fex1? fex2?* | This study |
| SDP 348 | *h+ mto2S338C:KanMX6 bgs1D277N Rlc1-tdTomato:NatMX6 Pcp1-mEGFP:KanMX6 ade6-M21x his3-Δ1 fex1Δ? fex2Δ? leu1-32 ura4-Δ18* | This study |
| SDP 362.1 | *h- mto1Δ:KanMX6 bgs1D277N Rlc1-tdTomato:NatMX6 Pcp1-GFP:KanMX6 ade6-M21x fex1? fex2? leu1-32 ura4-Δ18* | This study |
| SDP 366.1 | *h- bgs1D277N mto2[24A]:hphMX rlc1-tdTomato:NatMX6 pcp1-GFP:KanMX6 ade6-M21x leu1-32 ura4-Δ18 fex1? fex2?* | This study |
| SDP 373 | *h- mto2[24A]:hphMX rlc1-tdTomato:NatMX6 pcp1-GFP:KanR ade6-M210 leu1-32 ura4-Δ18* | This study |
| SDP 376 | *h+ Kan:pBgs1-mEGFP-Bgs1 Sfi1-mCherry:NatMX6 Rlc1-tdTomato:NatMX6 ade6-M21x leu1-32 ura4-Δ18* | This study |
| SDP 383 | *h- mto2S338N:KanMX6 KanMX6:Pbgs1-mEGFP-Bgs1 Sfi1-mCherry:NatMX6 Rlc1-tdTomato:NatMX6 ade6-M21x his3? leu1-32 ura4-Δ18* | This study |
| SDP 385.1 | *h- Sfi1-mCherry:NatMX6 Rlc1-tdTomato:NatMX6 GFP-Atb2:KanMX6 ade6-X leu1-32 ura4-Δ18* | This study |
| SDP 400.1 | *h- mto1Δ::kanMX6 Kan:pBgs1-mEGFP-Bgs1 Sfi1-mCherry:NatMX6 Rlc1-tdTomato:NatMX6 ade6-M21x leu1-32 ura4-Δ18* | This study |
| SDP 403 | *h+ Cdc7-mEGFP:KanMX6 Sfi1-mCherry:NatMX6 Rlc1-tdTomato:NatMX6 ade6-M21x leu1-32 ura4-Δ18* | This study |
| SDP 413 | *h- mto2S338N:KanMX6 Cdc7-mEGFP:KanMX6 Sfi1-mCherry:NatMX6 Rlc1-tdTomato:NatMX6 ade6-M21x his3? leu1-32 ura4-Δ18* | This study |
| SDP 421.1 | *h- leu1::kanMX6-P3nmt1-pkc1(HR1-C2)-mEGFP Rlc1-tdTomato:NatMX6 Pcp1-GFP:KanR ura4-Δ18 ade6-M210* | This study |
| SDP 429 | *h- mto2S338N:KanMX6 Sfi1-mCherry:NatMX6 Rlc1-tdTomato:NatMX6 GFP-Atb2:KanMX6 ade6-X leu1-32 ura4-Δ18* | This study |
| SDP 431 | *h- mto1Δ::KanMX6 Sfi1-mCherry:NatMX6 Rlc1-tdTomato:NatMX6 GFP-Atb2:KanMX6 ade6-X leu1-32 ura4-Δ18* | This study |
| SDP 432.1 | *h- mto2S338N:KanMX6 leu1::kanMX6-P3nmt1-pkc1(HR1-C2)-mEGFP Rlc1-tdTomato:NatMX6 Pcp1-mEGFP:KanMX6 ura4-Δ18 leu1-32 ade6-M21x* | This study |
| SDP 434.1 | *h- mto1Δ:KanMX6 leu1::kanMX6-P3nmt1-pkc1(HR1-C2)-mEGFP Rlc1-tdTomato:NatMX6 Pcp1-mEGFP:KanMX6 ade6-M21x leu1-32 ura4-Δ18* | This study |
| SDP 474 | *h+ mto1Δ::kanMX6 Cdc7-EGFP:KanMX6 Sfi1-mCherry:NatMX6 Rlc1-tdTomato:NatMX6 ade6-M21x leu1-32 ura4-Δ18* | This study |
| SDP 517.1 | *h- mto1(1-1085):ura4+ bgs1D277N Rlc1-tdTomato:NatMX6 Pcp1-GFP:KanMX6 ade6-M21x his3? leu1-32 ura4-Δ18 fex1? fex2?* | This study |
| SDP 522.1 | *h- mto1-427 bgs1D277N Rlc1-tdTomato:NatMX6 Pcp1-GFP:KanR ade6? fex1? fex2? his? leu1-32 ura4-Δ18* | This study |
